# Metagenomes from the Loxahatchee wildlife refuge in the Florida Everglades

**DOI:** 10.1101/2021.02.16.430518

**Authors:** David A Alvarez, Nikolya A Cadavid, Cale A Childs, Matthew F Cupelli, Victoria A De Leao, Alyssa M Diaz, Sophie A Eldridge, Yasmin B Elhabashy, Allison E Fleming, Nathan A Fox, Marianna Franco, James C Gaspari, Isabella M Gerstin, Kimberlee A Gibson, Alyssa L Huott, Alex O Johnson, Ellie G Majhess, Gabriela Mantilla, Gabriella S Perez, Juliet J Prieto, Bridget C Reutter, Elena I Rivera, Thomas R Rootes, Jade Sellers, Allison M Streibig, Joseph S Wilkinson, Siona Zayas-Bazan, Jehangir H. Bhadha, Alicia Clum, Christopher Daum, Tijana Glavina del Rio, Kathleen Lail, Simon Roux, Emiley A. Eloe-Fadrosh, Jonathan B. Benskin

## Abstract

The Florida Everglades ecosystem represents a significant wetlands area and serves as a terrestrial carbon reservoir mediated in large part by microorganisms. Shotgun metagenome sequencing provides a snapshot of microbial diversity and the frequency of metabolic and functional gene content. Here, we present an analysis of 20 sediment samples collected from the Arthur R. Marshall Loxahatchee National Wildlife Refuge to characterize the taxonomic and functional potential of the microbial and viral communities, and reconstructed metagenome-assembled genomes. A total of 122 medium-quality and 6 high-quality MAGs are reported, three of which likely represent a novel species within the class Dehalococcoidia.

The most abundant phyla of bacteria and archaea were Proteobacteria and Euryarchaeota, respectively. *Caudovirales* was the most abundant viral order. Significant differences in taxonomic composition and diversity were observed among collection sites. Additionally, water samples were analyzed for pH, total nitrogen, total organic carbon, elements (P, K, Mg, Fe, Mn, Pb, Ca, S), chloride, electric conductivity, orthophosphate, nitrate, and ammonia, while the sediment samples were analyzed for carbon, nitrogen, and pH. Differences in measured aquatic and sediment analytes revealed significant correlations with numerous phyla. Significant correlations were observed between estimated gene frequencies of both aquatic and sediment analytes, most notably between *kup/kdpB* and *dsrA/cysC* with potassium and sulfur, respectively, as well as *phoD/phnX* and *cysC* with pH. Together, these data provide an important view into the functional and metabolic potential encoded within the sediment microbial communities in the Florida Everglades.

## Background & Summary

The Arthur R. Marshall Loxahatchee National Wildlife Refuge encompasses 585 km^2^ of land in the Florida Everglades that has remained relatively undisturbed due to limited anthropogenic influences^1^. Agricultural fields dominate the Northern and Western sides of the refuge while the Eastern side is primarily bordered by urban development. The Southern portion is surrounded by a water conservation area (WCA) along with residential and agricultural land. Each of the four cardinal boundaries (North, South, East, and West) of the refuge has an inflow of water from their surrounding features, with a natural southbound flow. Along with local pump stations, Stormwater Treatment Areas (STA) 1E and 1W on both the Northeast and Northwest sides discharge into the flow of the L-40 and L-7 Canals which partially surround the refuge (Fig. 1a). Additionally, water from Lake Okeechobee passes through agricultural fields in the Everglades Agricultural Area (EAA) and then is discharged into the northern boundary of the refuge through multiple canals. On the western side of the refuge, water from Lake Okeechobee flows into the refuge through the Hillsboro Canal where it continues to flow to the Southern boundary. The Eastern and Northeastern boundaries of the refuge are predominantly residential, with neighborhoods approximately 120m away from the canal. Along with residential areas, the Eastern side of the refuge has boat rentals along with privately-owned agricultural fields bordering it, potentially allowing for greater anthropogenic influences on the refuge.

**Figure 1:**
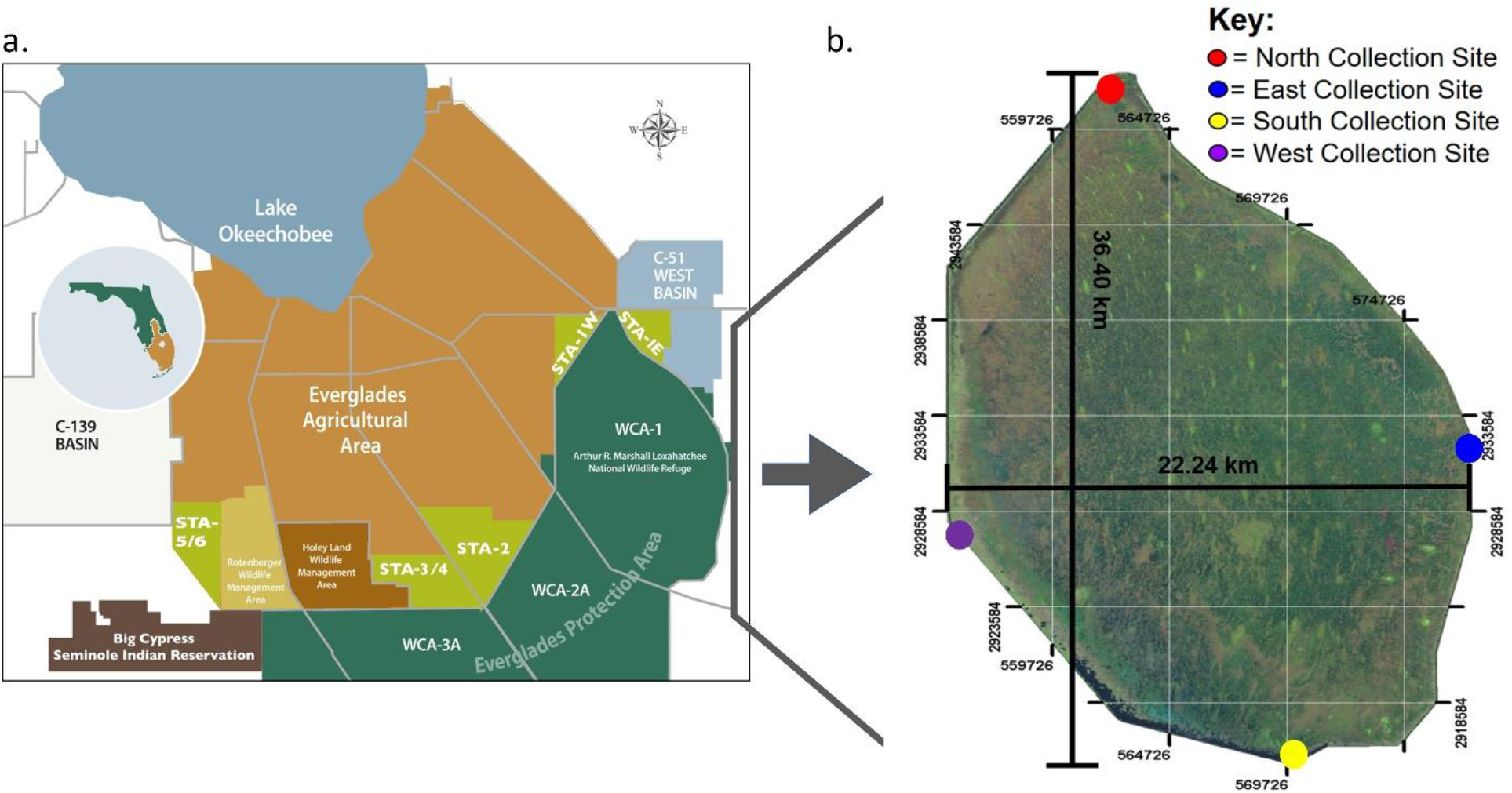
Sample location, sediment collection, and processing. **(a)** Stormwater Treatment Area map detailing waterflow and drainage stites through the EAA and Everglades Protection Area from the South Florida Water Management District (https://www.sfwmd.gov/document/stormwater-treatment-areas-map-O). (**b**) A display of the coordinate values of sample sites. North collection site: *26°40’38.8”N 80°22’31.2”W.* East collection site: *26°30’7.3”N 80°13’21.0”W.* South collection site: *26°28’08.5”N 80°26’36.4”W.* West collection site: *26°28’08.5”N 80°26’36.4”W.*

Previously, we applied shotgun metagenome sequencing to wetland soil samples collected across four sites within the Loxahatchee Refuge to investigate microbiome diversity and biogeochemically-relevant microbial processes^2^. In this report, we expand upon our initial study by collecting submerged sediment and water samples from the north, south, east and west sides of the refuge (Fig. 1b) to investigate (1) how location-specific environmental factors may contribute to differences in taxonomic diversity, (2) if potentially unique metagenome-assembled genomes (MAGs) can be assembled from sediment samples taken from the refuge, and (3) how environmental factors may impact the prevalence of genes related to energy cycling. Following sediment and water collection, DNA from the sediment samples was extracted, sequenced, assembled, and annotated. The sediment samples were analyzed for carbon, nitrogen, and pH, while the water samples were analyzed for pH, phosphorus (P), potassium (K), magnesium (Mg), iron (Fe), manganese (Mn), total nitrogen (TN), lead (Pb), electrical conductivity (EC), chlorine (Cl), carbon (C), ammonia, nitrate, sulfur (S), orthophosphates (Ortho-P), and calcium (Ca).

### Phylogenetic diversity

Environmental factors impact the taxonomic diversity of a sediment’s microbiome. This study aims to identify possible correlations between specific environmental factors and the microbial diversity found within aquatic sediment samples from the cardinal boundaries of the Loxahatchee Refuge.

Prokaryotic communities have been researched extensively in submerged aquatic sediment, and multiple environmental factors are known to strongly influence the taxonomic diversity of a sediment’s microbiome^3^. Notable variations in taxonomic diversity can occur with fluctuations in nitrogen concentration, organic carbon content, temperature, redox status, and pH^3^. Notably, a study from Chen^4^ demonstrated an increase in prokaryotic diversity near agriculturally dominated drainage sites. The addition of nitrogen to sediment has a direct effect on microbial communities and biomass because higher concentrations of nitrogen showed a decrease in sediment pH^5,6^ which can shift the taxonomic abundance of many bacterial phyla^7^. A negative correlation between nitrogen-based fertilizers and the abundance of Chloroflexi, Cyanobacteria, Firmicutes, and Planctomycetes has previously been reported^8^. Additionally, bacterial community richness and phylogenetic diversity increase at intermediate levels of salinity^9^.

While more is known about the diversity of aquatic viral communities, little is known about the diversity of viruses in submerged aquatic sediments. Potentially, the same biochemical factors impacting aquatic viral diversity will similarly affect viruses within the aquatic sediment. A review from Needham^10^ found that differences in the diversity of oceanic viral communities are impacted by nutrient availability, UV exposure, water temperature, and physiological properties of the virus and host. A preliminary study has shown that aquatic sediment from areas with differing levels of agricultural runoff do not have significant differences in viral diversity, implying that levels of agricultural runoff are not responsible for changes in viral taxonomic diversity^11^.

### MAGs

The second aim of our research is to assemble potentially novel MAGs to expand upon current knowledge of prokaryotic genomics and give public access to these datasets. Currently, no known MAGs have been published from Florida Everglades sediment. Past researchers have developed MAGs from other environments for purposes similar to the goals of this paper. In a study by Xue *et al*.^12^, prokaryotic MAG sequences were constructed from sediment samples to provide information on specific metabolic pathways and were reported in terms of completeness, contamination, size in base-pairs, GC content, and taxonomy^12^. MAGs were assembled and reported by similar standards in the present study.

### Energy Cycling

Our research also aims to address the estimated frequency of genes related to the environmental cycling of C, TN, P, S, and K (Table 1). The genes involved with nitrogen cycling that were analyzed include *narG,* which is utilized in nitrogen metabolism during dissimilatory nitrate reduction^13^. Furthermore, *nifH* is utilized in the fixation of atmospheric nitrogen into ammonia^14^; however, a negative correlation between nitrite and nitrate with *nifH* frequency has previously been observed^15^. The gene *cbbL* is a marker gene in the carbon cycle. Previously, a positive correlation between sediment organic carbon and *cbbL* abundance was observed^16^, making it possible that there could be a correlation between *cbbL* and TOC in the water column.

**Table 1:**
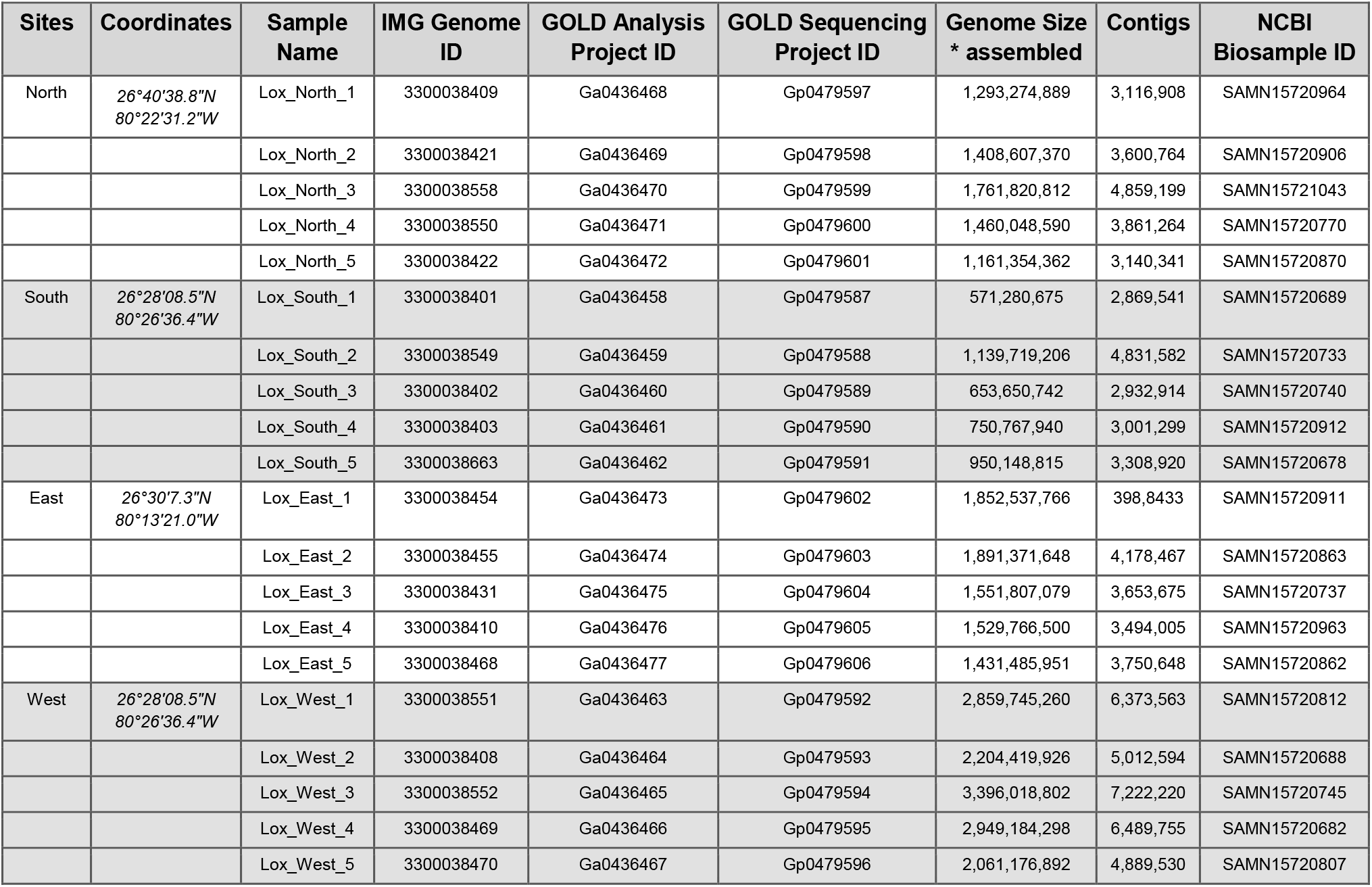
Details of twenty metagenomic samples taken from the Loxahatchee Wildlife Refuge.

Both *phoD* and *phnX* are involved with the hydrolysis of organic phosphates and phosphorus cycling. The *phoD* gene utilizes calcium ions as cofactors to hydrolyze a phosphate monoester and produce a free phosphate and alcohol^17^. Therefore, a positive correlation between the abundance of calcium and the frequency of *phoD,* and a negative correlation between orthophosphates and frequency of *phoD* is possible. Moreover, *phnX* produces phosphonoacetaldehyde hydrolase which hydrolyzes phosphonoacetaldehyde into a phosphate and an acetaldehyde^18^. Given the release of phosphates, a negative correlation between orthophosphates and the normalized gene values of *phnX* is possible. Moreover, as magnesium is a cofactor for phnX, a possible positive correlation between magnesium concentration and the normalized gene frequency of *phnX* might be observed.

The KDP-ATPase system can be affected by environmental conditions such as pH and potassium deficiencies in the extracellular environment^19^. The *kup* and *kdpB* genes within this system are responsible for potassium transport during cycling. Due to their role in potassium cycling, a positive correlation may exist between the frequency of *kdpB* and *kup* and the concentration of potassium in the samples.

The *dsrA* gene is involved in sulfate reduction^20^ and acts as a marker gene in microbes involved in sulfur cycling^21^, increasing the likelihood of a positive correlation between the frequency of *dsrA* and the concentration of sulfur in the water and the sediment. Additionally, it has been observed that *cysC* was positively correlated with sulfur concentrations in sediment^22^.

The *mcrA* gene is a marker gene involved in the methane metabolism pathway. A positive correlation between the nitrogen concentration and the gene frequency of *mcrA,* as well as a negative correlation with carbon concentration, have been observed in the wetland’s soil^2^. This suggests potential correlations between the abundance of the *mcrA* gene and the aquatic carbon and nitrogen concentrations.

Determining the estimated frequencies of these genes could allow conclusions to be drawn regarding possible anthropogenic influences on the microbiomes and their corresponding cycles. Furthermore, linking gene frequency to the concentration of certain factors at each site could reinforce findings seen in previous literature regarding the prevalence of certain genes in corresponding energy pathways.

## Results

Shotgun metagenomic sequencing of DNA from twenty sediment samples was performed using the Illumina NovaSeq S4 Sequencing system. A total of 3.29 x10^10^ base pairs (bp) were assembled (contigs ≥ 500bp) with an average length of 1.64 x10^8^ bp and 4.23 x10^6^ contigs per metagenome (Table 1).

Both the IMG pipeline^23^ and Kaiju^24^ were utilized for metagenome profiling, with IMG assigning taxa based on assembled data and Kaiju assigning taxa based on reads (Fig. 2). Only minor discrepancies were observed between these two methods (Supplementary Fig. S1). Proteobacteria was the most abundant bacterial phylum at each cardinal boundary with mean frequencies ranging from 0.518 (standard error of the mean (SEM) = 0.002) in the Southern samples to 0.546 (SEM = 0.003) in the Western samples (Fig. 2a). The most abundant generas were *Candidatus Solibacter* with a mean frequency of 0.017 (Eastern collection, SEM = <0.001), *Anaeromyxobacter* with a mean frequency of 0.016 (Northern collection, SEM = 0.002), and *Geobacter* with mean frequencies of 0.016 and 0.012, respectively (Western collection, SEM = <0.001 & Southern collection, SEM = 0.001) (Fig. 2d/e). These genera were present at all sites.

**Figure 2:**
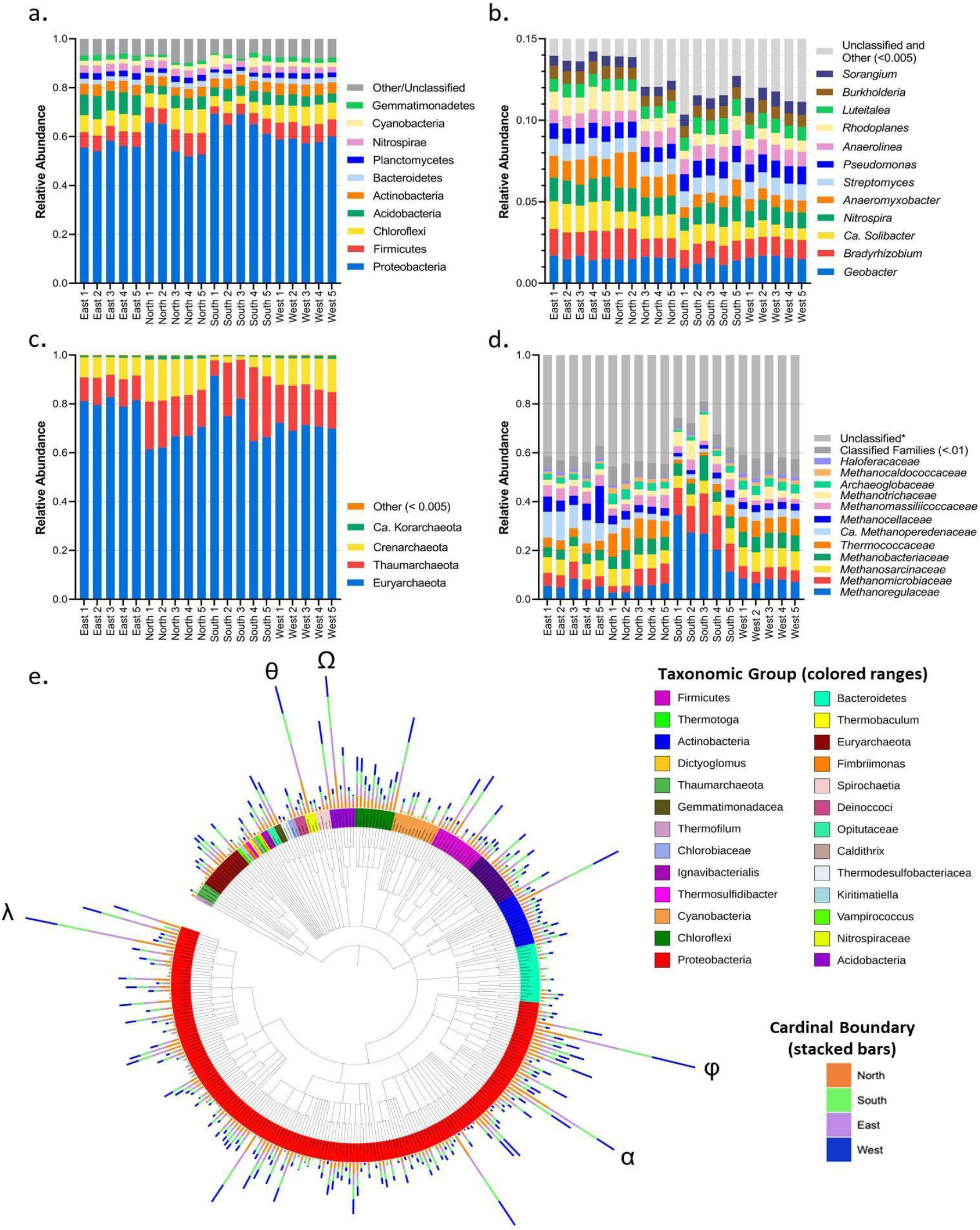
Taxonomic diversity and relative abundance from 20 metagenomic samples. Less abundant and unclassified were grouped together in panels a-d. (**a**) Relative frequency of the top-10 bacterial phyla present within all metagenomes as assigned by the IMG analysis pipeline. (**b**) Relative frequency of the top-12 bacterial genera present within all metagenomes as determined by Kaiju. (**c**) Relative frequency of the top-4 archaeal phyla present within all metagenomes. (**d**) Relative frequency of the top-12 archaeal families within phylum Eureoyarchaeota present within all metagenomes as assigned by the IMG pipeline. (**e**) Taxonomic groups (colored ranges) and genera (tree leaves) found within all metagenomic assemblies. Stacked bar charts surrounding the body of the tree compare the relative frequencies of genera from each of the cardinal boundaries. Any genus with a frequency <0.001 was not included. λ= *Bradyrhizobium,* α= *Anaeromyxobacter,* φ= *Geobacter,* Ω= *Candidatus Solibacter,* θ= *Nitrospira.*

Euryarchaeota was the most abundant archaeal phylum among all samples with the largest mean frequency (within archaea) located at the Eastern sites (0.808, SEM = 0.007) and lowest mean frequency located at the Northern sites (0.653, SEM = 0.016) (Fig. 2c). Within Euryarchaeota, the most abundant family was *Methanoregulaceae* with the largest mean frequency at the Southern site (0.240, SEM = 0.039) and lowest mean frequency at the Northern site (0.046, SEM = 0.008).

Simpson’s diversity (*D*), Shannon’s diversity *(He),* Pielou’s evenness (*Je*) and Margalef’s richness *(dMa)* indices were used to analyze alpha diversity (Fig. 3). All phylum diversity calculations reflect the data generated from both the IMG^23^ pipeline and Kaiju^24^. Bacterial phyla from the Eastern collection had the highest mean values of Simpson’s (*D*=0.664, SEM=0.007), Shannon’s (*He*=1.831, SEM=0.019) and Pielou’s (*Je*=0.457, SEM=0.005) indices. In contrast, bacterial phyla from the South showed the lowest mean values of Simpson’s (*D*=0.554, SEM=0.019), Shannon’s (*He*=1.563, SEM=0.052), and Pielou’s indices (*Je*=0.390, SEM=0.013). The Southern samples had the highest Margalef’s index mean (*dMa*=3.381, SEM=0.026) while the Western samples had the lowest Margalef’s index mean (*dMa*=3.126, SEM=0.018).

**Figure 3:**
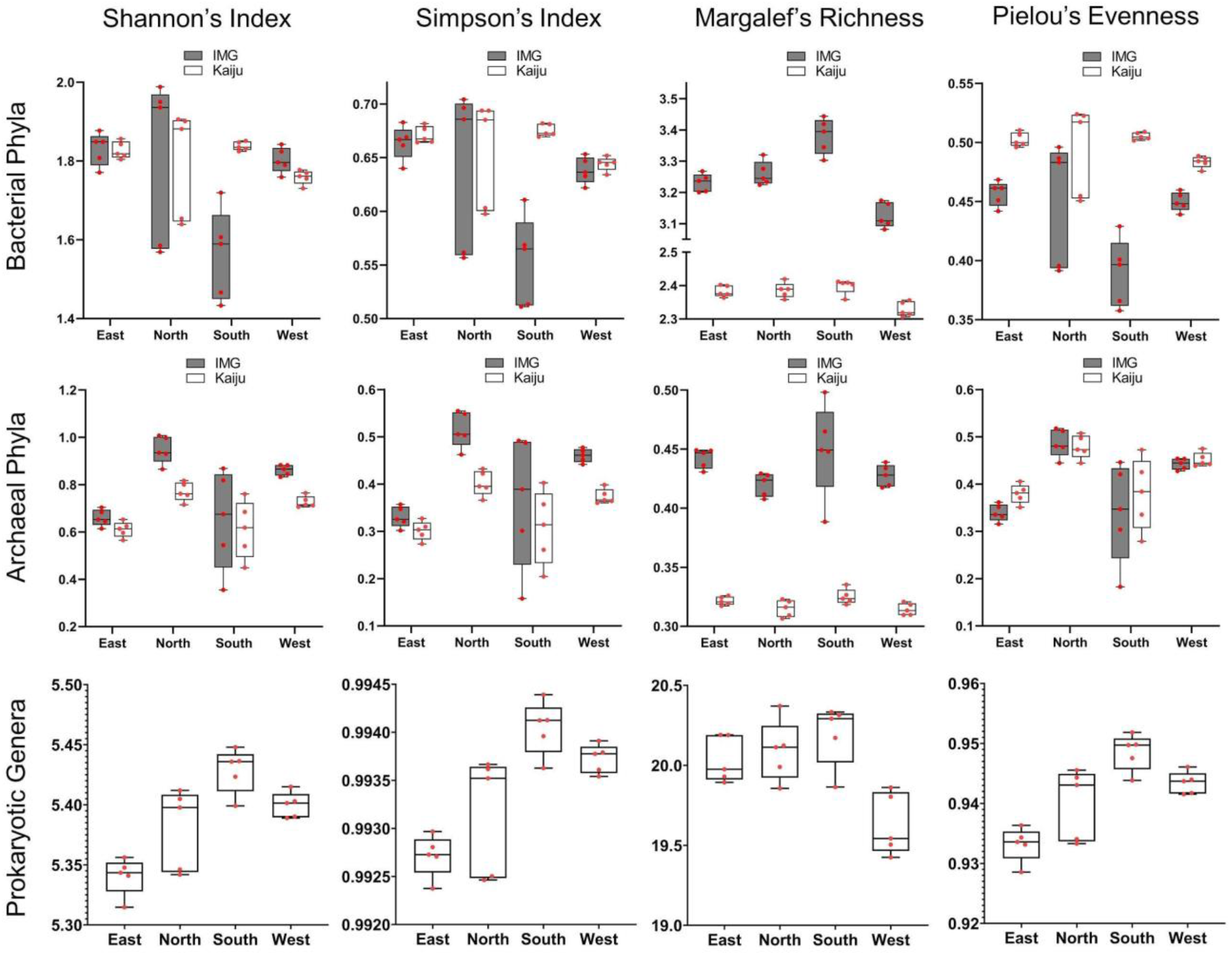
Matrix of Shannon’s index of diversity, Simpson’s index of diversity (dominance), Margalef’s richness index, and Pielou’s evenness index from the East, North, South, and West sites within Loxahatchee National Wildlife Refuge. Bacterial and archaeal phylum-level analysis completed utilizing data from the IMG (grey bars) and Kaiju (white bars) pipelines. Genus-level analysis was completed utilizing data from Kaiju. Rows represent bacterial phyla (top row), archaeal phyla (middle row) and prokaryotic genera (bottom row). The boxes depict the interquartile range (between the first and third quartiles), representing the middle 50% of the data. The horizontal line within the box represents the median while the whiskers indicate the lowest and highest values. Red points represent actual data values.

Archaeal phyla from the North collection displayed the highest e mean values for Simpson’s (*D*=0.515, SEM=0.017), Shannon’s (*He*=0.947, SEM=0.026), and Pielou’s (*Je*=0.487, SEM=0.013) compared to the other sites (Fig. 3). The sites with the lowest mean values for Simpson’s, Shannon’s, and Pielou’s indices were the South (*He*=0.652, SEM=0.093) and East (*D*=0.330, SEM=0.010, *Je*=0.339, SEM=0.008). The South site had the highest Margalef’s index mean (*dMa*=0.450, SEM=0.018) while the North site had the lowest (dMa=0.420, SEM=0.004).

Alpha diversity analyses were also conducted on prokaryotic genera with a frequency of >.001 (at one or more sites) using data generated by Kaiju^24^ (Fig. 3). The Southern samples had the highest mean Simpson’s (*D*=0.994, SEM=0.000), Shannon’s (*He*=5.429, SEM=0.008), Pielou’s *(Je=* 0.949, SEM=0.001), and Margalef’s (*dMa*=20.195, SEM=0.087) indices. Conversely, the Eastern collection showed the lowest means for Simpson’s (*D*=0.993, SEM=0.000), Shannon’s (*He*= 5.341, SEM=0.007), and Pielou’s (*Je*= 0.933, SEM=0.001) indices while the Western collection had the lowest mean for Margalef’s *(dMa=* 19.628, SEM=0.086).

Viral sequences were identified based on IMG^23^ taxonomic annotation. The number of unique viral strains identified in the metagenomic assemblies of the sediment samples ranged from 11 (South 1) to 65 (North 3) with means of 44.4 (SEM= 5.69), 30.4 (SEM= 4.83), 19.4 (SEM= 2.91), and 44.4 (SEM= 6.62) from the Northern, Southern, Eastern, and Western samples, respectively (Supplementary Fig. S2). The majority of identified viral genomes belonged to the order *Caudovirales,* with the exception of samples North 1 and 4, in which the predominant order was *Algavirales. Siphoviridae* was the most represented family across all samples within the order *Caudovirales,* followed by *Myoviridae* and *Podoviridae* (Fig. 4a). In North 1 and 3 along with East 2 and 3, *Phycodnaviridae* was the most represented family, falling within the order Algavirales (Fig. 4a). Importantly however, these represent only viruses identified based on similarity to reference viral genomes, while a large portion of the viral diversity remain entirely uncharacterized and could have gone undetected here^25^.

**Figure 4:**
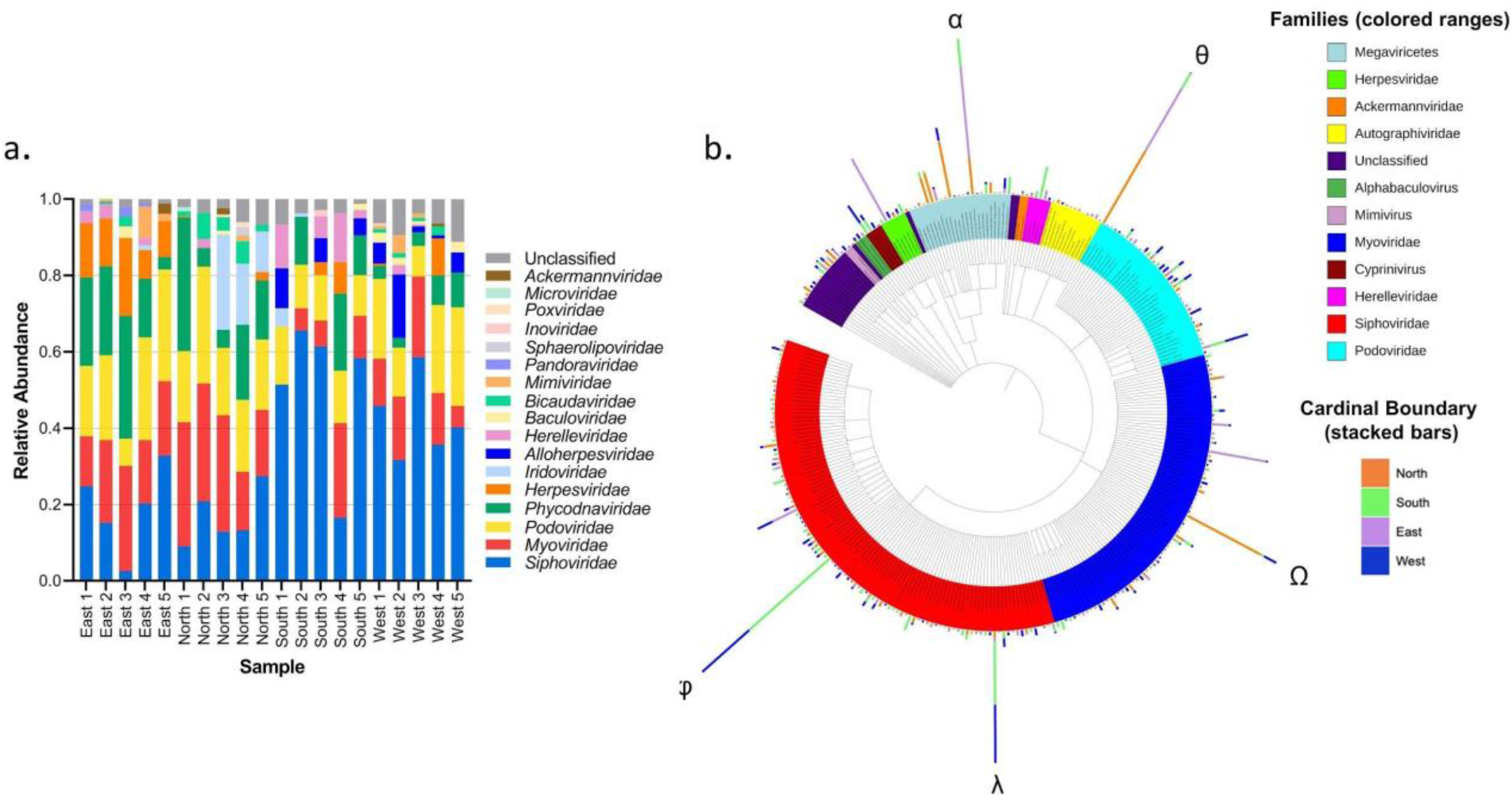
Taxonomic diversity and relative abundance of viral families from 20 metagenomic samples. **(a)** The relative abundance of each viral family present in each sediment sample as determined by the IMG pipeline. **(b)** Viral families (colored ranges) and strains (tree leaves) found within all metagenomic assemblies from the Loxahatchee Refuge. Stacked bars surrounding the body of the tree compare the relative frequencies of viral strains from each of the cardinal boundaries. φ= Actinoplanes phage phiAsp 2, λ= Pseudomonas virus NP1, Ω= Pseudomonas phage PhiPA3, θ=: Myxococcus phage Mx8, α= Bathycoccus sp. RCC1105 virus BpV.

Actinoplanes phage phiAsp 2 was the most abundant viral strain in the South collections with a mean relative frequency of 0.305 (SEM= 0.143), but was also present at a relatively high abundance in West samples with a mean frequency of 0.133 (SEM= 0.102) (Fig. 4b). This virus was not found in the North or East samples. Pseudomonas virus NP1 was the most abundant in the South collections, with a mean relative frequency value of 0.179 (SEM= 0.107), and was relatively low in abundance within the other sites. Both of these viruses belong to the family *Siphoviridae.* Pseudomonas phage PhiPA3 and Myxococcus phage Mx8 were both the most abundant in the North with mean frequency values of 0.141 (SEM=0.063) and 0.142 (SEM=0.037), respectively. Pseudomonas phage PhiPA3 was not as prevalent in any other sites. Myxococcus phage Mx8 was relatively abundant in the East, with a mean frequency of 0.125 (SEM=0.018), but not in the South or West samples. Bathycoccus sp. RCC1105 virus BpV was found to be the most abundant in the East, representing a mean frequency of 0.195 (SEM=0.027), but not present at high frequencies in any of the other locations.

Metagenomic binning using the IMG^23^ automated pipeline of the submerged sediment samples from the cardinal boundaries yielded 128 total bins: East (37), North (35), South (8), and West (48). Six of the bins were high quality (4.69%) and the other 122 bins were medium quality (95.31%) according to the minimum information about metagenome-assembled genomes (MIMAG) standards^26^. A table of MAGs can be found as Supplementary Table S1. Of the 128 total bins, 93 were bacterial and 35 were archaeal. The bins had a mean completeness of 69.02% (s.d.=12.89) and mean contamination of 3.16% (s.d.=2.44). The mean genome size was 2.265 Mb (s.d.=1.392 Mb). Additionally, read data was collected for each high quality (HQ) bin within all five metagenomic samples from the cardinal boundary which it originated from to assess if the MAG was ubiquitous within the cardinal boundary (Supplementary Table S3).

____Pairwise average nucleotide identity (ANI) was estimated between each HQ bin and its nearest neighbor on a phylogenetic species tree (see Methods). In each comparison, the ANI estimate was <80%, indicating that each HQ bin represents a separate operational taxonomic unit (OTU) from its known neighbors. Additionally, ANI estimates were determined between each pairing of HQ bins 3300038422_25, 3300038550_26, and 3300038558_31, as all were from the North sample site and had the same taxonomic neighbors. The ANI estimate from each of these comparisons was greater than 99.8%, indicating that these three MAGs most likely represent the same OTU. Phylogenetic trees and pangenomes were generated to display the ten closest neighbors for each HQ MAG (Fig. 5).

**Figure 5:**
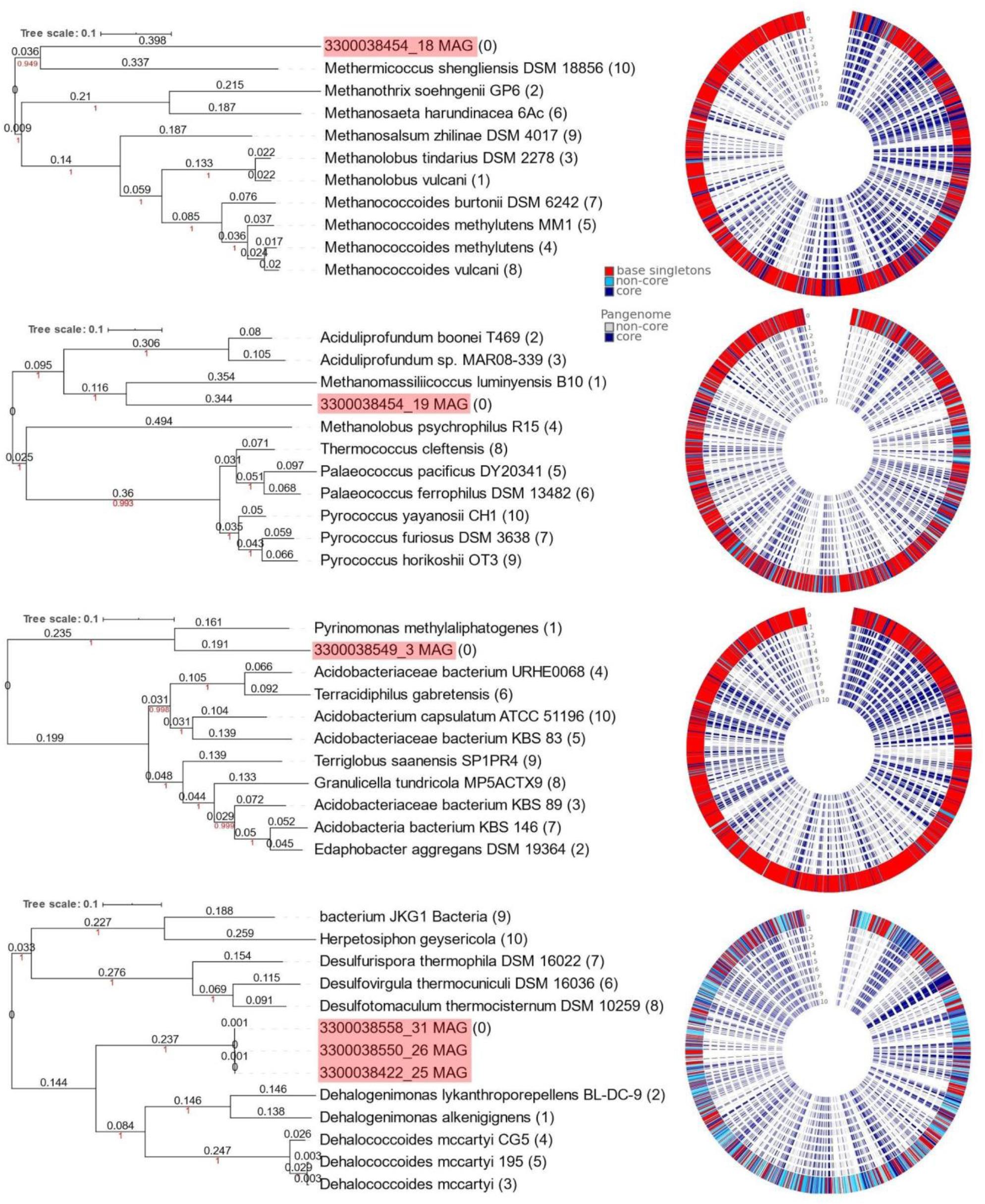
Taxonomic neighbors of all HQ MAGs. Trees display each HQ MAG and 10 of its closest neighbors. The black number displayed above the branches represents the branch length, red numbers represent the bootstrap value, and the numbers in parentheses represent the position in the accompanying pangenome. The outermost ring of each pangenome is the original HQ MAG at position 0 and the innermost ring is position 10. HQ MAGs 3300038422_25, 3300038550_26, and 3300038558_31 are collectively represented by 3300038558_31 on the accompanying pangenome because they represent the same OTU.

Functional analysis was performed using Clusters of Orthologous Groups (COGs). The COG categories in metagenomes from the same cardinal boundaries are ordinated closer to one another (Fig. 6a/b). Heirarcharcial clustering indicates considerable similarities between COG categories in the Eastern and Western samples (Fig. 6a), while PCoA indicates that COG categories in the East samples were more similar to the North samples (Fig. 6b). Most samples were relatively similar in the abundance of RNA processing and modification COGs (Fig. 6c).

**Figure 6:**
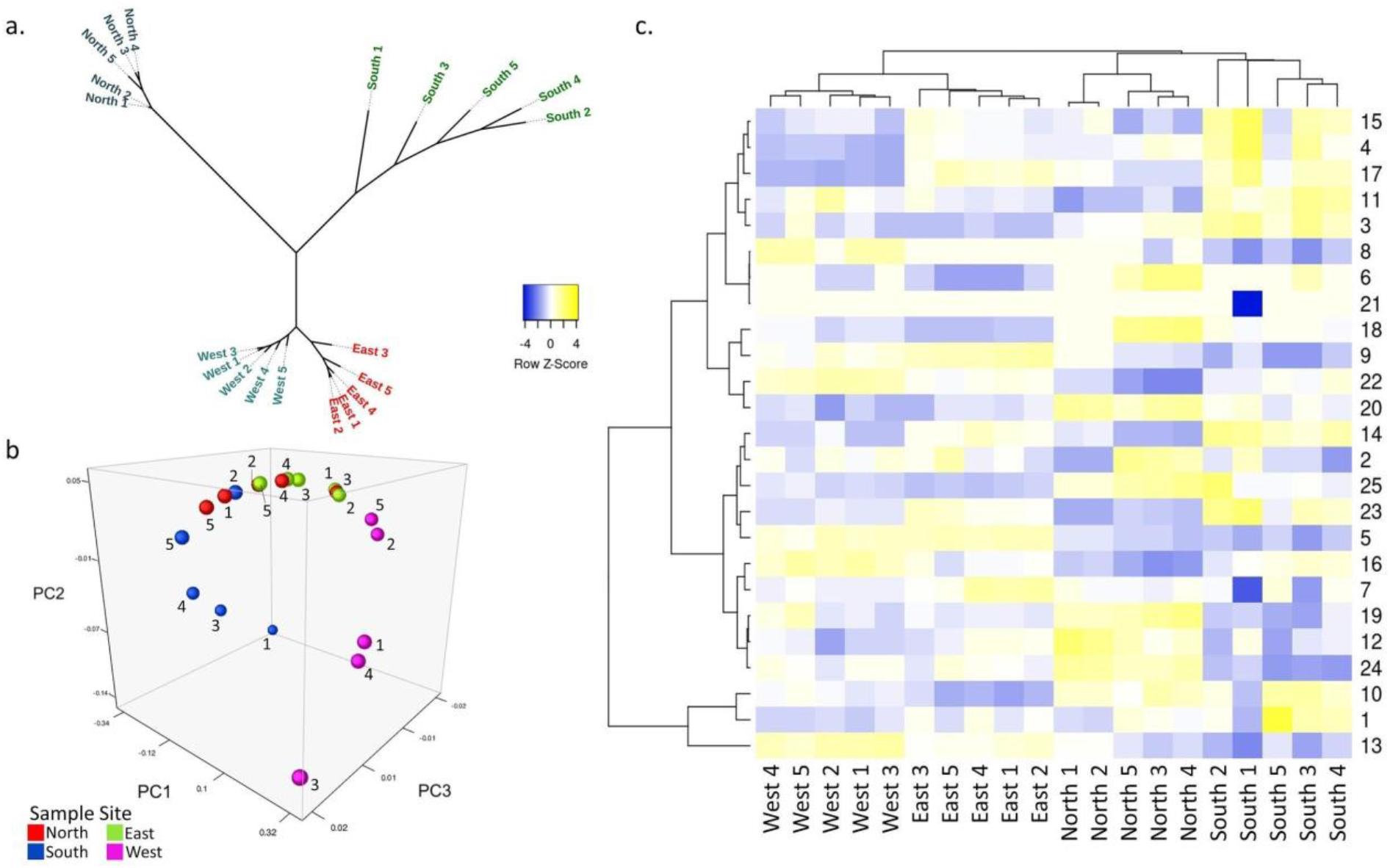
Functional analysis of metagenomic samples. (**a**) Unrooted tree displaying hierarchical clustering of COG categories. (**b**) PCoA of Genomic Clustering by COG category. (**c**) COG category cluster analysis based on the relative abundances of each metagenome’s protein dataset. The relative abundance of each COG category ranges from a low (blue) to a high (yellow). Different COG Categories: 1, Amino Acid Transport and Metabolism; 2, Carbohydrate Transport and Metabolism; 3, Cell cycle control, cell division, chromosome partitioning; 4, Cell Motility; 5, Cell wall/membrane/envelope biogenesis; 6, Chromatin structure and dynamics; 7, Coenzyme transport and metabolism; 8, Cytoskeleton; 9, Defense Mechanisms; 10, Energy production and conversion; 11, Extracellular structures; 12, Function unknown; 13, General function prediction only;14, Inorganic ion transport and metabolism;15,Intracellular trafficking, secretion, and vesicular transport; 16, Lipid transport and metabolism; 17, Mobilome:prophages, transposons;18, Nucleotide transport and metabolism;19, Post Translational modification, protein turnover; 20, Replication, recombination and repair; 21, RNA processing and modification; 22,Secondary metabolites biosynthesis, transport and catabolism; 23, Signal transduction mechanisms; 24, Transcription; and 25, Translation, ribosomal structure and biogenesis.

Three water samples were collected from each boundary and were analyzed to compare environmental factors to the relative frequency of genes involved with energy cycling. The three results from each aquatic analyte were averaged for each site in order to complete the Pearson’s rank correlation test against estimated gene frequencies. Nitrite and phosphorus were below the detection limit (<0.005 mg/L) in the water samples. Sediment from each metagenomic sample was analyzed for pH, C (%TOC), and N (%TN). A *p*-value <0.05 was considered statistically significant for this analysis. All results are found in Supplementary Table S2. Raw pH values are included in Supplementary Table S6.

### *nifH* and *narG*

A significant positive correlation was found between ammonia and *nifH (R=0.795, p*<0.001). Significant negative correlations were found between *nifH* and TOC (*R*=-0.481, *p*=0.032), K (*R*=-0.749, *p*<0.001), Mg (*R*=-0.737, *p*<0.001), EC (*R*=-0.633, *p*=0.003), S (*R*=-0.638, *p*=0.002), and Ca (*R*=-0.720, p<0.001). A significant positive correlation was found between ammonia and *narG (R=0.672, p*<0.001) as well as the sediment TOC content (*R*=-0.585, *p*=0.007). Significant negative correlations were observed between *narG* and K (*R*=-0.609, *p*= 0.004), Mg (*R*=-0.598 *p*= 0.005), EC (*R*=-0.644, *p*= 0.002), S (*R*=-0.598, *p*= 0.005), and Ca *(R=* −0.701, *p*=0.001).

### *kdpB* and *kup*

A significant negative relationship between *kdpB* and the concentrations of Ca (*R*= – 0.717, *p*<0.001), EC (*R*=-0.772, *p*<0.001), sediment TN (*R*=-0.57, *p=*0.009), sediment TOC (*R=*-0.639, *p*=0.002), and K (*R*=-0.782, *p*<0.001) was observed. The gene frequency of *kdpB* displayed a significant positive relationship with ammonia (*R*=0.569, *p*=0.009). Both aquatic pH (*R*=0.659, *p*=0.002) and ammonia (*R*=0.822, *p*<0.001) had a significant positive relationship with *kup* while a significant negative relationship was observed with K (*R*=-0.467, *p*=0.04), Mg (*R*=-0.446, *p*=0.05), sediment TN (*R=*-0.453, *p=*0.045), sediment TOC (*R=*-0.513, *p=*0.021), and Ca (*R*=-0.768, *p*<0.001). No significant correlations were observed between sediment pH and *kup/kdpB.*

### *dsrA* and *cysC*

There was a significant positive correlation between *dsrA* and aquatic (*R*=0.622, *p*=0.003) and sediment pH (*R*=0.543, *p*=0.02). A significant positive correlation between *cysC* and the concentration of S (*R=*0.528, *p*=0.02), Ca (*R*=0.637, *p*=0.003), sediment TOC (*R=*0.676, *p=*0.001), and sediment TN (*R=*0.650, *p=*0.002) was observed.

### *phoD* and *phnX*

A significant positive correlation was found between the normalized gene values of *phoD* and the sediment pH (*R*=0.484, *p*=0.042), aquatic pH (*R*=0.633, *p*=0.003), aquatic ortho-P (*R*=0.571, *p*=0.009) and Ca (*R*=-0.610, *p*=0.004). A significant negative correlation was found between the normalized gene values of *phnX* and the concentration of aquatic ortho-P (*R*=-0.730, *p*<0.001). Due to low phosphorus concentrations in the water samples, no analysis could be executed between normalized *phn*X values and phosphorus concentration.

### cbbL

A positive correlation between the abundance of *cbbL* and the concentration of ammonia (*R*=0.765, *p*<0.001) and aquatic pH (*R*=0.540, *p*=0.01) was observed. A negative relationship was found between *cbbL* and Ca (*R*=-0.540, *p*=0.01).

### mcrA

There was a positive relationship between *mcrA* and the concentration of ammonia (*R*=0.834, *p*<0.001). A negative relationship was found between *mcrA* and K (*R*=-0.701, *p*<0.001), Mg (*R*=-0.686, *p*<0.001), EC (*R=*-0.653, *p=*0.002), S (*R*=-0.619, *p*=0.004), and Ca (*R*=-0.765, *p*<0.001).

### PCoA

PCoA was performed utilizing only the functions/KOs included in this study (Supplementary Fig. S3). The PCoA showing all ten functions/KOs showed clear clustering according to sample sites. PCoA utilizing only *nifh/narG, phoD/phnX,* or *dsrA/cysC* shows clustering of the North and East sites, while *kdpB/kup* shows a standardized clustering of all sites except the South site.

## Discussion

The most abundant bacterial phyla found within all samples were Proteobacteria, Firmicutes, and Chloroflexi (Fig. 2a). Proteobacteria are found in most freshwater environments, as they are able to withstand an array of different temperatures and pressures, emphasizing their role as one of the most versatile and abundant bacterial phyla^27^. Certain Proteobacteria have symbiotic relationships with plant roots, possibly contributing to their high frequencies within these sediment samples^28^. A previous metagenomic study conducted on the eastern side of the interior of the refuge found that the most abundant phyla in submerged sediment samples were Proteobacteria, Acidobacteria, and Actinobacteria^2^, while our current study found the three most abundant phyla were Proteobacteria, Firmicutes, and Actinobacteria. The most common archaeal phyla found within the sediment samples were Euryarchaeota, Thaumarchaeota, and Crenarchaeota. The phylum Euryarchaeota has remained largely unexplored, contributing to the lack of knowledge regarding the biogeographic role of these organisms in terrestrial ecosystems^29^. However, a study of the biological diversity within subtropical coastal wetland sediments observed that the archaeal communities identified consisted primarily of organisms belonging to Euryarchaeota^30^, agreeing with the results of this study.

Alpha diversity results show that the South site consistently had lower diversity and evenness in phyla while the North site showed higher diversity and evenness (Fig. 3 and Supplementary Table S10).The southern site was the only site that consisted of mostly sandy sediment (the other sites had less sand were a mud-like consistency), which may have contributed to the diversity and evenness of the prokaryotes within. The North site would likely contain agricultural runoff (the EAA and STA-1W is located just to the North of the collection site) which corroborates with results from Chen^4^. In contrast to the phylum-level analyses, alpha diversity analyses of genera revealed that the South site showed the highest mean values in all indices, while the East site displayed the lowest.

Previous studies noted that conditions such as vegetation and historic temperature between the topsoil and subsoil layers can influence the bacterial and archaeal diversity in collections^31,32^. Although precautions were taken to ensure consistent sampling methods, minor differences in the depth of the cores could have contributed to the differences between the sites.

When comparing the alpha diversity results obtained from the IMG^23^ pipeline versus Kaiju^24^, the results were relatively similar, with the exception being Margalef’s index and results from the southern site (Fig. 3). Kaiju^24^ assessed the richness to be lower than IMG^23^ for both the bacterial and archaeal phyla due to 44.7% more bacterial phyla and 40% more archaeal phyla being identified by IMG^23^. In addition, there was a discrepancy between the South results (bacterial phyla) for Simpson’s, Shannon’s, and Pielou’s indices potentially due to the differences in computational methods utilized to measure abundance by Kaiju^24^ and IMG^23^ (Supplementary Table S10).

Spearman’s analysis showed significant correlations between alpha diversity indexes and water/sediment analytes (Supplementary Table S9a and S9b). For the bacteria, a negative correlation between pH and Simpson’s (aquatic: *p*=0.002, ρ= −0.65; sediment: *p*=0.002, ρ=-0.675), Shannon’s (aquatic: *p*=0.006, ρ= −0.589; sediment: *p*=0.008, ρ=-0.606), and Pielou’s (aquatic: *p*=0.006, ρ=-0.589; sediment: *p*=0.008, ρ=-0.606) indexes was observed, indicating that pH has a significant impact on the bacterial microbiome. The pH level could be a contributing factor to lower phylum diversity at the southern site, as it displayed the highest mean aquatic and sediment pH. Potassium (*p*= 0.000, ρ= −0.822), magnesium (*p*=0.000, ρ= −0.822), and sulfur (*p*=0.000, ρ= −0.822) concentrations all demonstrated a negative correlation with the bacterial Margalef’s index which may have contributed to the western site’s lower richness.

In contrast to the bacteria, the archaeal alpha diversity was not significantly impacted by pH. Instead, significant correlations were observed for both sediment TOC (Simpson’s *p*=0.012., ρ=0.552; Shannon’s *p*=0.001, ρ=0.669; Margalef’s *p*=0.022, ρ=-0.508; Pielou’s *p*=0.001, ρ=0.669) and sediment TN (Simpson’s *p*=0.026, ρ=0.496; Shannon’s *p*=0.003, ρ=0.627; Margalef’s *p*=0.012, ρ=-0.551; Pielou’s *p*=0.003, ρ=0.627) which supports that these key nutrients impact archaeal diversity.

Significant correlations between phylum abundance and pH were found for Acidobacteria (aquatic: *R*= −0.633, *p*=0.003, sediment: *R*= −0.860, *p*= 0.000), Armatimonadetes (aquatic: *R*= −0.815, *p*=0.000, sediment: *R*= −0.666, *p*=0.003), Chrysiogenetes (aquatic: *R*= 0.514, *p*=0.021, sediment: *R*=0.698, *p*=0.001), Nitrospirae (aquatic: *R*= −0.629, *p*=0.003, sediment: *R*= −0.630, *p*=0.005), Planctomycetes (aquatic: *R*= −0.583, *p*=0.007, sediment: *R*= −0.744, *p*=0.004), and Spirochaetes (aquatic: *R*= 0.532, *p*=0.016, sediment: *R*=0.617, *p*=0.006). These results corroborate with previous findings from Bartram *et al.*^7^. Numerous other significant correlations between observed phyla and environmental analytes were found. Aquatic S, EC, Ca, Mg, K, and ammonia in addition to sediment TOC and TN significantly impacted the largest number of phyla (Supplementary Fig. S4).

Sediment samples obtained from the South boundary of the refuge exhibited decreased viral diversity and richness in comparison to the East, North, and West samples (Supplementary Fig. S2). Our finding of lowered viral diversity in sandy substrates agrees with previous studies^33^. The low viral abundance in the South could also be attributed to the small amount of DNA that was assembled from the samples (Supplementary Fig. S2). A positive correlation between the phosphate concentration and the rate of viral production has been documented in oceanic surface waters^34^. Our study was unable to replicate these findings, as the South samples had the highest mean concentration of ortho-P but the lowest number of identified viral strains (Supplementary Fig. S2). However, it is important to note that viral communities collected by Motegi *et al.^34^* were extracted from seawater, while the viral communities in this study were extracted from sediment.

Viruses belonging to the family *Siphoviridae* constituted less than 20% of all viruses present in the North and East samples, but 54.6% of the viruses present in the samples obtained from the West and 50.6% of the viruses from the South. One possible explanation for this observation is the decreased frequency of Proteobacteria in the North and East samples, in addition to the increased frequency of Proteobacteria in the West samples (Fig. 2a). Further, the mean relative frequency of *Siphoviridae* was highest within the South sediment samples (Fig. 4a) potentially due to the increased frequency of *Methanomicrobiaceae* (which is preyed upon by *Siphoviridae)* in these samples (Fig. 2d). It has been observed that the relative abundance of *Siphoviridae* correlates with the relative abundance of *Methanomicrobia^35^,* the class in which *Methanomicrobiaceae* belongs.

Five viral strains emerged in overwhelmingly abundant frequencies. However, these results represent the relative viral abundances of our sediment samples and may not accurately represent the actual viral abundances of the sampling sites. *Pseudomonas virus NP1* was one of the five most abundant viruses, found in large frequencies in the South and East sediment samples (Fig. B). *Bathycoccus sp. RCC1105 virus BpV* is a marine virus, with close relatives previously observed in freshwater samples, and was found here in abundance primarily at the North and East collection sites (Fig. B).

Although bacteriophages are the most abundant type of virus, their diversity and abundance is difficult to estimate due in part to an absence of standardized metrics^36^; however, metagenomics has been exceedingly helpful in quantifying their abundance. *Actinoplanes phage phiAsp 2* was one of the most abundant phages in the West and South samples, yet not in the North or East (Fig. B). Phage *phiAsp 2* infects the bacterial genus *Actinoplanes,* which is typically found in sediment^37^. Thus, the presence of phage *phiAsp2* in our samples was not unexpected. *Myxococcus phage Mx8* targets the bacterium *Myxococcus xanthus^38^* which has an optimum pH range of 5-8^39^, matching the pH ranges of both the sediment and water samples of this study. *Pseudomonas phage phiPA3* was found in only the North samples (Fig. B) and infects the bacteria *Pseudomonas aeruginosa.* These bacteria live in a wide range of environments^40^ (Monson, 2011) and ecological niches^41^, making it plausible to be found in the refuge.

### MAGs

Metagenome assembly and binning resulted in 128 MAGs ranging across 21 different phyla. Differences between the most abundant taxa identified in the samples and the constructed MAGs show that MAGs are not necessarily representative of the taxonomy in each sample; however, eight of the ten most abundant bacterial phyla (Nitrospirota, Planctomycetota, Bacteroidota, Actinobacteriota, Acidobacteriota, Chloroflexota, Firmicutes, and Proteobacteria) and one of the four most abundant archaeal phyla across the samples (Crenarchaeota) were represented as constructed MAGs.

A Pearson’s rank correlation test found a positive correlation between metagenome size and the number of MAGs assembled from each sample (*R*=0.779, *p*<.001). This positive correlation may explain the lower number of MAGs generated from Southern samples, as these samples had the smallest genome sizes and the smallest quantity of MAGs of any site.

The results of mapping each HQ bin against all of the metagenomic samples within their respective sites revealed that each displayed >0% mapped reads in at least four out of the five metagenomic samples (Supplementary Table S3), suggesting that each HQ MAG is ubiquitous at the cardinal boundary it originated from. ANI analysis revealed that each of the HQ bins represents a different OTU than its closest neighbor; however, these neighbors give the best insight into the biological relevance of these organisms at their respective sample sites.

Among the assembled MAGs from the South 5 sample was a bin of the class *Lokiarchaeia* (bin ID: 3300038663_10), which is part of the Asgard group of organisms. These organisms (Lokiarchaeota, in particular) have provided insight into the emergence of eukaryotes from prokaryotes^42^. Lokiarchaeia has not previously been reported in the Florida Everglades ecosystem; therefore, this constructed MAG from the South sample site represents a novel location for the presence of this group of archaeal organisms.

The HQ bin 3300038454_18 from the East 1 metagenome was identified as an OTU within the Archaeal class Methanosarcina. The closest neighbor to this bin as identified by the species tree is *Methermicoccus shengliensis DSM 18856.* This anaerobic species has previously been identified in the Shengli oilfield in China. The species is known to be thermophilic and can grow at temperatures around 65°C. Additionally, *M. shengiensis* is methanotrophic and was found to have increased growth with the addition of sludge^43^.

The other HQ bin from the East 1 metagenome (3300038454_19) was assigned to the archaeal class Thermoplasmata_A. The nearest taxonomic neighbor observed on the species tree is *Methanomassiliicoccus luminyensis B10,* a methanogen that produces methane gas via the reduction of methanol and oxidation of hydrogen.

The HQ bin 3300038549_3 from South 2 was identified to be from the class Blastocatellia. The species tree revealed the bin’s closest neighbor to be *Pyrinomonas methylaliphatogenes,* an aerobic, heterotrophic, moderately acidophilic, and thermophilic bacterium. It has been proposed that *P. methylaliphatogenes* scavenges for hydrogen when there is a lack of organic electron donors, therefore contributing to hydrogen cycling^44^.

The three HQ MAGs from the North sample sites (3300038422_25, 3300038550_26, and 3300038558_31) were shown to be part of the same OTU following the results of ANI analysis (all three had ANI >99%). This indicates the presence of a prominent organism of the bacterial class Dehalococcoidia that is unique to the North site, as it could not be mapped to any of the other sample sites in notable amounts. Also, the separate assembly of this OTU from multiple metagenomic samples (North 3, 4, and 5) indicates that it is unlikely that this MAG was developed as a result of an assembly error. The closest taxonomic neighbors to this OTU were *Dehalogenimonas lykanthroporepellens, Dehalogenimonas alkenigignens,* and *Dehalococcoides mccartyi.* These bacterial species are relevant to the degradation of chlorinated organics in contaminated environments via reductive dechlorination or dehalogenation^45^. However, a preliminary analysis indicated these MAGs do not contain the reductive dehalogenase markers TIGR02486 or pfam13486.

### Functional Analysis and Energy Cycling

The North samples appeared to be the most distinct in context of COG categories (Fig. 6). This dissimilarity might be attributed to runoff entering the North site from STA^46^, Lake Okeechobee, residential areas, and the EAA. Although significant correlations were observed between estimated gene frequency and water parameters, it should be stated that water movement, inconsistency in the water column, and change over time will gradually alter sediment composition and the microbiome within. Also, the correlations stated in this section are based upon estimated gene frequency rather than gene activity.

The results showed a positive correlation between *nifH* and ammonia (*R*=0.795, *p*<0.001), possibly due to the activity of nitrogen fixating genes which can increase ammonia levels in the water^47^. Of the studied genes, *nifH* had the smallest normalized values, which may be a result from the lack of direct contact with atmospheric nitrogen utilized by *nifH.* Further, the positive correlation between *narG* and ammonia (*R*=0.672, *p*=0.001) could be due to *narG’s* role in dissimilatory nitrate reduction^13^. Despite the presence of iron-sulfate clusters in *narG*^48^, our results showed a negative correlation between sulfur (*R*=-0.598, *p*=0.005) and *narG.* This may be due to the measurement accounting for environmental sulfur in the water and not the sediment.

The positive correlations between normalized values of *phoD* and aquatic (*R=*0.633 *p=0.003)* and sediment (*R*=0.484, *p*=0.042) pH are supported by previous findings that pH is a significant environmental factor in bacterial expression of the *phoD* gene^49^. We anticipated a positive correlation between calcium and *phoD,* similar to findings that *phoD* activity was enhanced through calcium supplementation^50^; however, our results demonstrated a negative correlation between *phoD* and Ca (*R*=-0.610 *p*=0.004). This difference may be explained by the previous study^50^ utilizing recombinant *phoD* genes whereas our study examined wildtype *phoD.* The negative correlation between the concentration of ortho-P in the water column and *phnX* (*R*=-0.730 *p*<0.001) could be due to a decreased demand for organic phosphate decomposition while in the presence of ortho-P, as the gene is activated in conditions of phosphate deficiency^51^. This assertion, however, is challenged by the positive correlation between *phoD* and ortho-P concentrations (*R*=0.571 *p*=0.009). As *phoD* has a similar role in organic phosphate decomposition to *phnX,* we expected the genes to have complementary relationships with ortho-P.

A significant positive correlation was found between *dsrA* and pH of both the water (*R*=0.622, *p*=0.003) and the sediment (*R*=0.543, *p*=0.02), but a direct causal relationship could not be elucidated from this study. A positive correlation was observed between *cysC* and aquatic sulfur concentration (*R=*0.528, *p*=0.02), possibly due to the role of *cysC* in reducing sulfate to sulfide^22^.

A significant negative correlation was found between *kdpB* abundance and potassium concentration (*R*=-0.782, *p*<0.001). This may be due to *kdpB’s* key role in potassium cycling, as a low concentration of available potassium would lead to more organisms needing to uptake as much potassium as possible^52^. Our results found a significant positive correlation between *kup* abundance and the pH of the water column (*R*=0.659, *p*=0.002), however, a significant correlation was not observed with the sediment pH. A negative correlation was observed between *kup* and potassium (*R*=-0.467, *p*=0.04) potentially due to water nutrient levels not representing the nutrient levels in the sediment where the gene abundance was derived. Furthermore, a significant negative correlation was observed between *kup* and Ca (*R*=-0.768, *p*<0.001), in contrast to previous findings that Ca sensors could mechanistically link abiotic stress in plants and the microbiome to several K+ uptake systems including *kup*^53,54^.

The positive correlation between *cbbL* and ammonia (*R*=0.765, *p*<0.001) may be attributed to the high abundance of *cbbL* in wetland environments. Wetlands tend to have higher ammonia concentrations due to a lack of oxygen preventing nitrogen conversion^55^. The positive correlation between *mcrA* and ammonia (*R*=0.834, *p*<0.001) is potentially due to ammonia acting as an inhibitor of McrA^56^. The positive correlation between *mcrA* and K (*R*=-0.701, *p*<0.001) may be because available K is a significant influence on methanogenic communities, including those involving *mcrA^57^.*

PCoA analysis of the *kup/kdpB* and *dsrA/cysC* gene pairs found strong similarities in clustering when comparing to the PCoA analysis of all ten genes involved in this study (Supplementary Fig. S3). It is possible that potassium and sulfur are important elemental factors driving the functional profile of this environment. Significant correlations in the frequency of several genes and the pH of both the sediment/water were observed, suggesting that pH is also an important factor.

### Limitations

Multiple methods were utilized to obtain taxonomic classifications. Although discrepancies were present, each method demonstrated similar results. Both methods resulted in approximately 50% of the sample remaining unidentified, indicating further studies are needed to truly examine the taxonomic diversity within sediment samples. There is an evident need for consistent, in-depth computational methods to more clearly explore and study the phage/viral abundance, distribution, and gene coverage in natural ecosystems. With the construction of well-established methods and metrics, novel phage characterizations and behavioral descriptions are likely to result. Additionally, it was not possible to fully characterize the alpha diversity at the genus-level due to computational limitations that prevented the identification of the genera found in frequencies less than 0.01%.

Due to the COVID-19 pandemic, the laboratory space that contained the sediment samples was inaccessible. After DNA extraction, these sediment samples were left for a period of time while drying, possibly altering the quantities of C, N, and pH present in the samples before analysis. Additionally, the Pearson’s correlation test relied on analytes from the water above the sediment samples and thus may not accurately reflect the levels within the sediment at the exact time of collection. Furthermore, the fluid nature of the environment makes replication of this experiment difficult.

While the MAG trees and analysis were comprehensive, they do not provide a full picture of the ecosystem the samples were gathered in. In order to obtain a full picture of the environment, the three sections of this paper must be examined in conjuncture.

## Conclusion

The microbiome results reported within this study have greatly expanded upon the current collection of metagenomic data from the Everglades ecosystem. The analyses within this study have determined both consistencies and distinctions between the taxonomic diversity of the microbiomes at each of the four cardinal boundary sample sites of the Loxahatchee refuge. A number of specific, potentially novel microorganisms were also identified via MAG assembly.

Significant correlations were found between the water/sediment analytes and the taxonomic abundance, alpha diversity and gene frequency of the microbiomes. Genes related to sulfur and potassium cycling were the most similar to the overall PCoA analysis conducted, and therefore have a large impact on the energy cycling processes in the environment while pH, %TN, and %TOC had significant impacts on overall microbiome diversity.

Observable differences in the microbiomes of the refuge canals will continue to experience longterm changes due to past, present, and future anthropogenic influences. Future research is required to expand upon the taxonomic diversity, gene abundance, and the prokaryotic genomic profile of the Everglades microbiome. The taxonomic profiles reported in this study can serve as a basis for further research conducted on the Everglades ecosystem.

## Methods

### Sediment Collection, DNA Extraction, and Water/Sediment Analytical Analysis

All samples were collected from the Arthur R. Marshall Loxahatchee National Wildlife Refuge in Palm Beach County, Florida, on February 2, 2020. Sediment was collected using a coring device from a depth of ~0.6m. Five samples were collected from each of the cardinal boundaries 20-40 cm apart (twenty samples total). All samples were taken from the inner bank of the canal except Lox North 1, taken on the outer bank of the refuge. Three water samples (~1.5L) were taken from each site. Each site had similar weather, wind speed, and air temperature (Supplementary Table S7). The North collection site had primarily tall, grass-like vegetation with a few trees growing on the banks. The East site had an abundance of tall vegetation and thick reeds, but was much denser and more difficult to find shallow water suitable for collecting sediment samples. The South and West collection sites were similar with dense patches of reeds, and flora visible just beneath the surface. Sediment from the North, East, and West sites were similar with a thick, mud-like consistency and abundant plant matter, while sediment from the South had a large amount of sand/shells but less plant material (differences could be due to the construction of three impoundment areas for water management purposes by the Army Corps of Engineers in the late 1950s, consisting of a series of levees and canals^58^.

Sediment samples were sieved through 2mm sterilized mesh. The recommended protocol for the QIAGEN DNeasy^®^ PowerSoil^®^ Kit (QIAGEN, Hilden, Germany) for DNA extraction was utilized for all samples on the same day as the collection. Frozen/thawed sediment samples from the West site underwent an additional, modified extraction process two weeks after collection due to low DNA content during initial extraction. In this modified procedure, samples containing 1.5x the amount of sediment were centrifuged for twice as long and bead-beated for 1.7x longer than recommended.

All water analyses (pH, TN, TOC, elements P, K, Mg, Fe, Mn, Pb, Ca, S, chloride, electric conductivity, ortho-phosphate, nitrate, and ammonia) were completed at the University of Florida/IFAS Everglades Research and Education Center (Belle Glade, FL). A Fisher Scientific Accument AB250 pH meter (Waltham, United States) was used for pH measurement. Standard pH buffers (20mL; pH 4, 7 and 10) were used for calibration. Ten mL of each water sample was analyzed until a constant pH was recorded. A Shimadzu (Kyoto, Japan) TOC-L analyzer was utilized (TNOC method) for TN and TOC measurement. The TN & TOC standards, quality control (QC) standards, and 40ml of each water sample were utilized in the analysis. ICP-OES was used for testing each element using the EPA 200.7 method^59^. A colorimeter equipped with 15 mm tubular flow cell and 480 nm filters was used to measure chloride concentration using the EPA 325.2 method^60^. A conductivity bridge, range 1 to 1000 μmho per centimeter, 5.2 conductivity cell, cell constant 1.0 or micro dipping type cell with 1.0 constant YSI #3403 or equivalent, and a 5.3 thermometer were used to measure electrical conductivity with the EPA 120.1 method^61^. An automated continuous flow analyzer was used for the analysis of ortho-phosphate concentration, nitrate concentration, and ammonia. Ortho-phosphate concentration was measured following the EPA 365.1 method^62^. Nitrate (NO3-N) with cadmium reduction was completed utilizing the EPA 353.3 method^63^. Ammonia (NH3-N) was measured utilizing the EPA 350.1 method^64^.

All sediment analyses (pH, %TOC, %TN) were completed at the Stable Isotopes for Biosphere Science Laboratory, Department of Ecology and Conservation Biology, Texas A&M University. Sifted sediment samples were dried to a constant weight and ground to fine powder using Retesch Oscillating Mixer Mill MM400 (Haan, Germany). Soil pH was determined by using an Accumet Basic pH meter (Denver Instrument, Arvada, CO, USA) on a 1:2 solution of soil in a 0.01 M CaCl2 solution^65^. The samples were analyzed for %TOC and %TN using the Costech Elemental Combustion System (Costech Analytical Technologies, Santa Clarita, CA, USA) coupled to a Thermo Conflo IV Interface (Thermo Fisher Scientific, Waltham, MA, USA) and a Thermo Scientific Delta V Advantage Stable Isotope Mass Spectrometer (Thermo Fisher Scientific, Waltham, MA, USA). The NIST plant standard Apple1515 was used to calculate the Nitrogen and Carbon concentrations (%). Carbon isotope ratios and %TOC were measured using acid-treated samples. Samples weighed in silver capsules were left with hydrochloric fume in a dessicator for 8 hours. After they are tested for residual carbonate, they were dried at 60C for 12 hours. Full results of the analysis are found in Supplementary Table S6a and S6b.

The mean value of each aquatic analyte (n=3) and sediment analyte (n=5, with exceptions for the pH-see Supplementary Table 6Sb) was utilized for the analysis of each cardinal boundary.

### Sequencing, Quality Control, Assembly, and Annotation

The Illumina NovaSeq 6000 platform (Illumina, San Diego, CA, USA) was used for metagenome sequencing using the S4 Regular flowcells and generated 6.24E+09 filtered reads with 9.05E+11 filtered bp. BBDuk (version 38.79)^66^ was used to trim and remove reads. Reads mapped with BBMap^66^ to contaminants were separated. The final filtered fastq files were error corrected using BBCms (version 38.44)^66^. The readset was then assembled using metaSPAdes (version 3.13.0)^67^. The input read set was mapped to the final assembly and coverage information generated with BBMap (version 38.44)^66^.

BBDuk (version 38.79)^66^ was used to remove contaminants, trim reads that contained adapter sequences, right quality trim reads where quality drops to 0, and remove reads that contained 4 or more ‘N’ bases, had an average quality score across the read less than 3, or had a minimum length <= 51 bp or 33% of the full read length. Reads mapped with BBMap^66^ to masked human, cat, dog, and mouse references at 93% identity were separated into a chaff file, as well as reads aligned to common microbial contaminants. The final filtered fastq files were error corrected using BBCms (version 38.44)^66^ using the command line options mincount=2 and highcountfraction=0.6.The readset was then assembled using metaSPAdes (version 3.13.0)^67^ using the following command line options: --only-assembler -k 33,55,77,99,127 --meta. The input read set was mapped to the final assembly and coverage information generated with BBmap (version 38.44)^66^ using the following command line options: bbmap.sh ambiguous=random.

Assembled metagenomes were processed through the DOE-JGI Metagenome Annotation Pipeline and loaded into the Integrated Microbial Genome & Microbiomes platform (IMG/M)^23^. Sample metadata is available through the Genomes OnLine Database (GOLD)^68^.

### Taxonomy

Taxonomic data was obtained with the JGI IMG portal^23^. The total amount of estimated gene copies of the phyla and families were counted for each metagenome. The normalized (estimated) frequency of each was found by dividing the estimated phylum/family count for a specific classification by the total estimated count for all the phyla/families from the given metagenome. Additionally, Kaiju (version 1.1.2)^24^ on KBase^69^ was also utilized to establish taxonomic assignments using the unassembled, filtered metagenomes.

Using data derived from Kaiju^24^ and IMG^23^, newick tree files were generated for phyla/families/genera using the NCBI Taxonomy Browser^70,71^ found at https://www.ncbi.nlm.nih.gov/Taxonomy/CommonTree/wwwcmt.cgi. Repeats were eliminated. The resulting taxonomic cladograms were visualized using iTOL (version 4)^72^. For each tree, the abundance of each taxonomic assignment was represented by compiling the datasets for the individual samples at the four cardinal boundaries and entered utilizing the iTOL Annotation Editor^72^.

Simpson’s Index D, Shannon’s Index, Pielou’s Index, and Margalef’s Index were calculated with the phylogenetic distribution of genes from the IMG portal^23^ utilizing “by BLAST percent identities” as well as phylogenetic estimates from Kaiju (version 1.1.2)^24^.

Genome bins were identified using the metagenome binning tool, MetaBAT^73^. A consensus lineage was determined per bin using the IMG taxonomy of individual contigs, and contigs with a different phylum assignment than the consensus were removed. GTDBTK lineage was also assigned using GTDB-tk (version 0.2.2) with the GTDB database (release 86)^74^. Genome completeness and contamination were estimated using CheckM (version 1.0.12) based on the recovery of core single-copy marker genes^75^. The bins were sorted into High, Medium, and Low quality and reported according to the Genomic Standards Consortium (GSC)-compliant Minimum Information on a Metagenome-Assembled Genome (MIMAG)^26^.

Individual high quality MAG trees were constructed utilizing KBase^69^. The high quality MAG assemblies were downloaded from IMG^23^ and then run through the app “Annotate Microbial Assembly”^76^ to convert the FASTA file into a genome file. The genome was then run through the app “Insert Genome into Species Tree-v2.2.0”^77^ utilizing the following settings: “Neighbor Public Genome Count” = 10 and “Copy Public Genomes to Workspace” = yes. The resulting newick file was then uploaded to iTOL (version 4) and edited in the iTOL annotation editor^72^.

Pearson’s correlation tests were used to analyze relationships between the normalized frequencies of each phyla assigned from the IMG pipeline and average concentration of each analyte in the water and actual values for the sediment samples. Spearman’s correlation tests were utilized to analyze the relationships between the taxonomic abundance of phyla along with alpha diversity means against the average concentration of each analyte in the water and actual values for the sediment samples. These statistical analyses along with all stacked bar and box/whisker charts were performed/created by Graphpad Prism version 8.4.2 for Windows (GraphPad Software, San Diego, California USA, www.graphpad.com).

### MAGs

The JGI IMG pipeline^23^ was utilized to generate MAGs from the sequenced data. Genome bins were identified with the automated metagenome binning tool, MetaBAT (version 2.12.1)^73^. Each MAG was manually exported from the JGI IMG portal^23^ and uploaded to KBase^69^ to construct phylogenetic trees and uploaded to iTOL (version 4)^72^.

_____A pairwise ANI calculation was performed between each HQ bin and its nearest taxonomic neighbor as visualized in the species tree output using the KBase^69^ application “Compute ANI with FastANI” with default settings^78^. The KBase application “Align Reads using Bowtie2-v2.3.2” was used to determine the raw count and percent of mapped reads for each genome bin^79^. All settings were “default”. Pangenomes for each HQ MAG were created utilizing the genome sets created when running “Insert Genome into Species Tree” (version 2.2.0)^80^. These genome sets were then used to calculate the pangenome utilizing “Build Pangenome with OrthoMCL” (version 2.0)^81^ and then visualized utilizing “Pangenome Circle Plot” (version 1.2.0)^81^ with all settings “default”.

### Functional Analysis and Energy Cycling

The genomes of all samples were clustered using the JGI IMG database “Compare Genomes’“ function under “Genome Clustering” via hierarchical clustering and grouped by COG categories^23^. The data generated was exported as a newick file, uploaded into iTOL (version 4)^72^, and displayed as an unrooted tree (Fig. 6a). Metagenomes were analyzed using the JGI IMG database “Compare Genomes” function, comparison via PCoA^23^. Data was exported to Altius Institute’s ‘CubeMaker’ (https://tools.altiusinstitute.org/cubemaker/#) to generate Fig. 6b and Supplementary Fig. S3. Data used to generate the COG category heatmap (Fig. 6c) were obtained on the JGI website by examining the overview of each genome^23^. A table of estimated gene counts and percent total of the different COG categories was recorded for each metagenome. The percent total of each specific COG category was then uploaded to Heatmapper Expression with average linkage and separated using the Euclidean distance method (http://www.heatmapper.ca/expression/)^82^.

Each gene included in this study was identified from the sample sites using their KEGG Orthology (KO), and the estimated abundance of each gene (>30% identity) was found by matching genes with reference genes on the IMG platform^23^. Normalized gene values were derived by dividing the number of estimated gene copies for each gene from each sample site by the total number of genes assembled in each metagenome and were used as gene frequencies. Pearson’s correlation tests were used to analyze relationships between the gene frequencies and average concentration of each analyte in the water and sediment samples.

## Acknowledgements

The authors would like to thank the Joint Genome Institute for their ongoing, generous support and help. The work conducted by the U.S. Department of Energy Joint Genome Institute, a DOE Office of Science User Facility, is supported by the Office of Science of the U.S. Department of Energy under Contract No. DE-AC02-05CH11231. We would like to thank the Sid Kyle Endowed Chair in Arid and Semiarid Land Biogeochemistry from the Department of Ecology and Conservation Biology (Texas A&M University) for sediment analysis as well as Dr. Ayumi Hyodo, Dr. Ulrich Stingl (UF), Dr. Willm Martens-Habbena (UF) and Ivica Letunic (iTOL) for their support and advice. Additionally, we would like to thank Dr. Abul Rabbany and the University of Florida-Soil, Water, and Nutrient Management Lab in Belle Glade, Florida for facilitating a portion of the analytical work. Lastly, we would like to thank the Loxahatchee National Wildlife Refuge for granting us access to collect water and sediment samples.

## Author Contributions

### Boca Raton Community High School

DAA: Energy cycling and sediment collection. NAC: Energy cycling and Sediment collection. CAC: MAGs and Field Photography. MFC: Metadata collection and taxonomic analysis. VAD: Water sample collection and beta diversity analysis. AMD: Site photographer and beta diversity analysis. SAE: Collection logistics and viral taxonomy. YBE: DNA extraction, sterilization, and alpha diversity analysis. AEF: Metadata collection and energy cycling. NAF: Energy cycling and collection navigation. MF: Sterilization, water testing methods, water analysis and functional analysis. JCG: Energy cycling and sediment collection. IMG: Collection logistics and viral taxonomy. KAG: DNA extraction and taxonomy. ALH: DNA extraction and energy cycling. AOJ: Water sample collection and MAGs. EGM: Sediment collection, metagenomic sequencing results, viral taxonomy, and functional analyses. GM: Sediment collection and taxonomy. GSP: DNA extraction and MAGs. JJP: Collection logistics and taxonomy. BCR: Collection logistics and energy cycling. EIR: DNA extraction and taxonomy. TRR: Sediment collection and MAGs. AMS: DNA extraction and taxonomic diversity analysis. JSW: MAGs, Sample collection, and DNA extraction. SZB: MAGs and collection quality control.

### Everglades Research & Education Center, University of Florida

JB: Analytical water analysis.

### Joint Genome Institute, U.S. Department of Energy

KL: Library preparation and QC (Quality Control). CD: Sequencing. ACl: Metagenome assembly and QC. ACo: Metagenome assembly and QC. RS: Data analysis. TGdR: Coordinated sequencing and analysis. EAE-F: supervised sequencing and analysis.

All questions and correspondence should be sent to Jon Benskin.

