## Supplementary Materials for "Metagenomes from the Loxahatchee wildlife refuge in the Florida Everglades"

Alvarez, David A[1†]- Energy cycling and sediment collection
Cadavid, Nikolya A[1†]- Energy cycling and sediment collection
Childs, Cale A[1†] - MAGs and Field Photography
Cupelli, Matthew F[1†]- Metadata collection and taxonomic analysis
De Leao, Victoria A[1†]-Water sample collection and beta diversity analysis
Diaz, Alyssa M[1†]- Site photographer and beta diversity analysis
Eldridge, Sophie A[1†]- Collection logistics and viral taxonomy
Elhabashy, Yasmin B[1†]- DNA extraction, sterilization, and alpha diversity analysis
Fleming, Allison E[1†]- Metadata collection and energy cycling
Fox, Nathan A[1†]- Energy cycling and collection
Franco, Marianna[1†]- Sterilization, water testing methods, water analysis and functional analysis
Gaspari, James C[1†]- Energy cycling and sediment collection
Gerstin, Isabella M[1†]- Collection logistics and viral taxonomy
Gibson, Kimberlee A[1†]- DNA extraction and taxonomy
Huott, Alyssa L[1†]- DNA extraction and energy cycling
Johnson, Alex O[1†]- Water sample collection and MAGs
Majhess, Ellie G[1†]- Sediment collection, metagenomic sequencing results, viral taxonomy, and functional analysis
Mantilla, Gabriela[1†]- Sediment collection and taxonomy
Perez, Gabriella S[1†]- DNA extraction and MAGs
Prieto, Juliet J[1†]- Collection logistics and taxonomy
Reutter, Bridget C[1†]- Collection logistics and energy cycling
Rivera, Elena I[1†]- DNA extraction and taxonomy
Rootes, Thomas R[1†]- Sediment collection and MAGs
Sellers, Jade[1†] - Sediment collection and taxonomy
Streibig, Allison M[1†]- DNA extraction and taxonomic diversity analysis
Wilkinson, Joseph S[1†]- MAGs, Sample collection, and DNA extraction
Zayas-Bazan, Siona[1†]- MAGs and collection quality control
Bhadha, Jehangir H.[2]- Water analysis
Clum, Alicia[3]- Metagenome assembly and QC
Daum, Christopher[3]- Sequencing
Glavina del Rio, Tijana[3]- Coordinated sequencing and analysis
Lail, Kathleen[3]- Library preparation and QC (Quality Control)
Roux, Simon[3]- Data analysis and manuscript preparation
Eloe-Fadrosh, Emiley A.[3]- Supervised sequencing and analysis
Benskin, Jonathan B.[3]- Corresponding author

**Affiliations:**
[†]Contributed equally to this work
[1]Boca Raton Community High School
[2]Everglades Research & Education Center, University of Florida
[3]Joint Genome Institute, U.S. Department of Energy

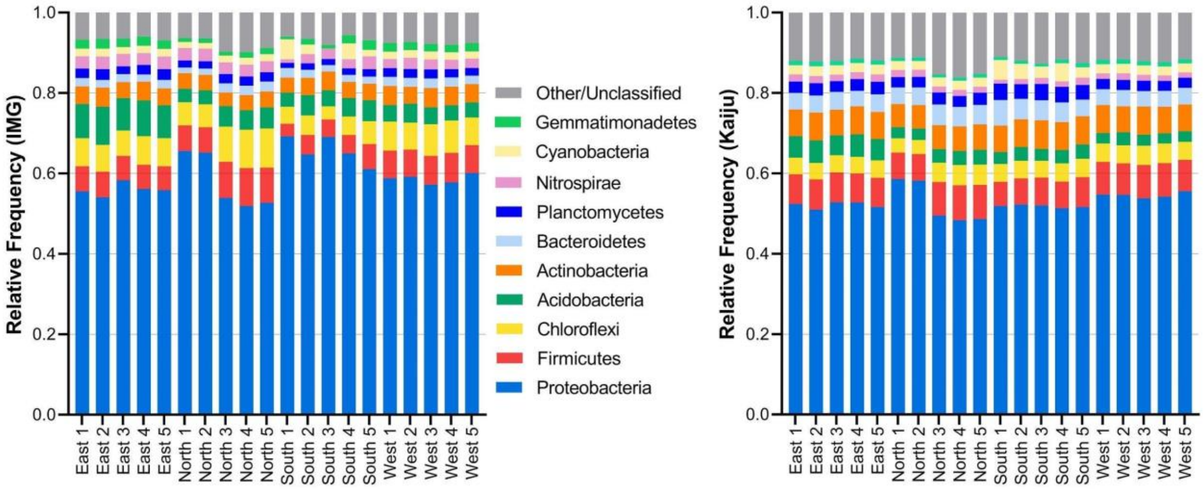

**Supplementary Figure S1:** Comparison of IMG and Kaiju binning methods for the top-10 phyla found within all twenty metagenomic samples. Only minor differences between these methods/results were observed.

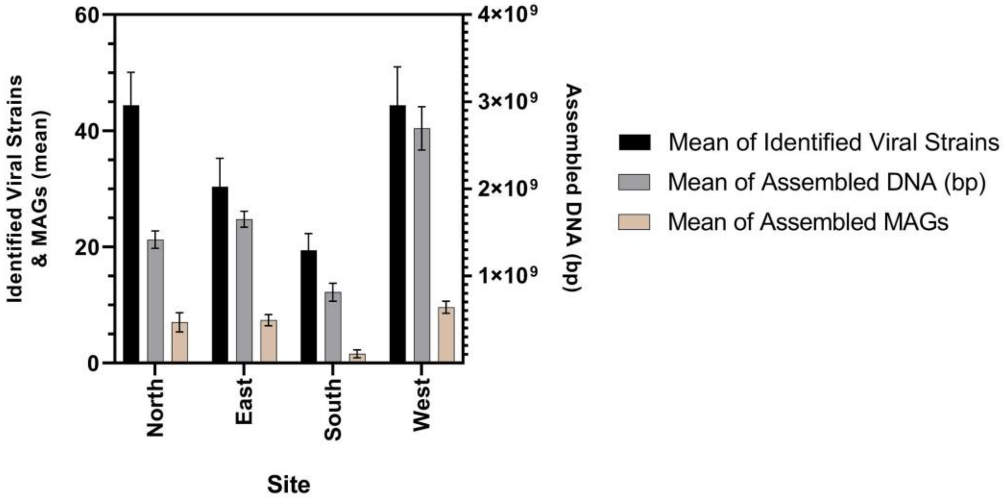

**Supplementary Figure S2:** Means of viral genotype richness, abundance, and assembled MAGS for each cardinal boundary from the five samples collected at each cardinal boundary.. Error bars = SEM.

**Supplementary Table S1:** All HQ and MQ MAGs with >50% completion and <10% contamination. Taxonomy, percent completeness and contamination, genome size, scaffold and gene count, MIMAG status, 16s rRNA counts, mapped read, and mean coverage data of bins.

| Site | Bin ID | Phylum | Taxonomy | Completeness | Contamination | Genome Size (bp) | Scaffold Count | Gene Count | MIMAG classification | 16S | Mean Coverage | Mapped Reads (%) | Number of Mapped Reads |
|---|---|---|---|---|---|---|---|---|---|---|---|---|---|
| East | 3300038410_12 | Chloroflexota | Bacteria; Chloroflexota; Anaerolineae; SBR1031; A4b | 55.17 | 0.91 | 3,997,766 | 637 | 4109 | MQ | 0 | 8.53 | 0.09 | 228,071 |
| East | 3300038410_16 | Chloroflexota | Bacteria; Chloroflexota; Ellin6529; CSP1-4; CSP1-4 | 54.63 | 2.01 | 1,977,389 | 201 | 1990 | MQ | 0 | 15.12 | 0.08 | 199,827 |
| East | 3300038410_17 | Halobacterota | Archaea; Halobacterota; Methanosarcinia | 85.4 | 4.25 | 1,833,504 | 189 | 2072 | MQ | 1 | 11.24 | 0.06 | 137,702 |
| East | 3300038410_20 | Nitrospirota | Bacteria; Nitrospirota; Nitrospiria; 2-01-FULL-66-17 | 62.27 | 2.12 | 1,522,042 | 149 | 1555 | MQ | 1 | 14.60 | 0.06 | 148,500 |
| East | 3300038410_21 | Nitrospirota | Bacteria; Nitrospirota; Thermodesulfovibrionia; Thermodesulfovibrionales; SM23-35 | 50.87 | 0 | 1,175,393 | 238 | 1411 | MQ | 0 | 9.78 | 0.03 | 76,824 |
| East | 3300038410_22 | Halobacterota | Archaea; Halobacterota; Methanosarcinia; Methanocellales; Methanocellaceae; Methanocella | 58.2 | 1.14 | 1,140,477 | 248 | 1434 | MQ | 0 | 10.12 | 0.03 | 77,211 |
| East | 3300038410_8 | Chloroflexota | Bacteria; Chloroflexota; Anaerolineae; Thermoflexales | 77.43 | 9.4 | 5,516,427 | 822 | 5401 | MQ | 2 | 11.71 | 0.17 | 431,800 |
| East | 3300038410_9 | Chloroflexota | Bacteria; Chloroflexota; Anaerolineae; Anaerolineales; envOPS12; UBA12294 | 79.8 | 4.08 | 5,476,203 | 785 | 5801 | MQ | 1 | 8.67 | 0.13 | 317,211 |
| East | 3300038431_11 | Chloroflexota | Bacteria; Chloroflexota; Anaerolineae; Anaerolineales; envOPS12; UBA12294 | 72.19 | 4.73 | 4,700,176 | 823 | 5033 | MQ | 0 | 7.89 | 0.08 | 247,699 |
| East | 3300038431_12 | UBP1 | Bacteria; UBP1; PRR-12 | 90.34 | 9.89 | 3,378,216 | 366 | 3096 | MQ | 0 | 9.87 | 0.08 | 222,743 |
| East | 3300038431_14 | Chloroflexota | Bacteria; Chloroflexota; Ellin6529; CSP1-4; CSP1-4 | 71.45 | 3.01 | 2,267,406 | 268 | 2334 | MQ | 0 | 20.89 | 0.11 | 316,332 |
| East | 3300038431_15 | Nitrospirota | Bacteria; Nitrospirota; Nitrospiria; 2-01-FULL-66-17 | 71.72 | 3.91 | 1,961,951 | 170 | 2016 | MQ | 0 | 17.19 | 0.08 | 225,264 |
| East | 3300038431_18 | Halobacterota | Archaea; Halobacterota; Methanomicrobia; Methanomicrobiales | 91.72 | 0.65 | 1,604,728 | 176 | 1932 | MQ | 0 | 8.26 | 0.03 | 88,558 |
| East | 3300038431_20 | Crenarchaeota | Archaea; Crenarchaeota; Bathyarchaeia; 40CM-2-53-6; RBG-16-50-20; RBG-16-50-20 | 68.09 | 1.46 | 1,236,319 | 217 | 1515 | MQ | 0 | 10.84 | 0.03 | 89,508 |
| East | 3300038431_21 | Thermoplasmatota | Archaea; Thermoplasmatota; Thermoplasmata_A; RBG-16-68-12; RBG-16-68-12 | 63.27 | 3.49 | 1,064,042 | 157 | 1235 | MQ | 0 | 55.40 | 0.13 | 393,734 |
| East | 3300038454_11 | Chloroflexota | Bacteria; Chloroflexota; Anaerolineae; Anaerolineales; envOPS12; UBA12294 | 81.64 | 2.25 | 6,420,605 | 808 | 6670 | MQ | 1 | 9.53 | 0.14 | 408,578 |
| East | 3300038454_14 | Chloroflexota | Bacteria; Chloroflexota; Anaerolineae; SBR1031; A4b | 65.51 | 3.18 | 4,322,233 | 602 | 4412 | MQ | 0 | 9.48 | 0.09 | 273,785 |
| East | 3300038454_15 | UBP1 | Bacteria; UBP1; PRR-12 | 77.89 | 5.28 | 2,807,266 | 425 | 2727 | MQ | 0 | 9.85 | 0.06 | 184,588 |
| East | 3300038454_18 | Halobacterota | Archaea; Halobacterota; Methanosarcinia | 96 | 3.27 | 2,164,102 | 201 | 2414 | HQ | 2 | 11.74 | 0.06 | 169,697 |
| East | 3300038454_19 | Thermoplasmatota | Archaea; Thermoplasmatota; Thermoplasmata_A; RBG-16-68-12; RBG-16-68-12 | 91.39 | 3.2 | 1,992,330 | 164 | 2220 | HQ | 1 | 81.58 | 0.36 | 1,085,636 |
| East | 3300038454_21 | Halobacterota | Archaea; Halobacterota; Methanosarcinia; Methanocellales; Methanocellaceae; Methanocella | 77.62 | 0.98 | 1,807,369 | 285 | 2148 | MQ | 1 | 12.89 | 0.05 | 155,791 |
| East | 3300038454_22 | Nitrospirota | Bacteria; Nitrospirota; Nitrospiria; 2-01-FULL-66-17 | 73.18 | 2.32 | 1,808,060 | 125 | 1829 | MQ | 0 | 17.21 | 0.07 | 207,887 |
| East | 3300038454_24 | Nitrospirota | Bacteria; Nitrospirota; Thermodesulfovibrionia; Thermodesulfovibrionales; SM23-35 | 69.82 | 2.73 | 1,641,482 | 240 | 1877 | MQ | 0 | 12.80 | 0.05 | 140,352 |
| East | 3300038455_18 | Chloroflexota | Bacteria; Chloroflexota; Anaerolineae; Anaerolineales; UBA4823 | 57.64 | 2.81 | 2,660,757 | 513 | 2793 | MQ | 1 | 11.57 | 0.06 | 205,544 |
| East | 3300038455_19 | Thermoplasmatota | Archaea; Thermoplasmatota; Thermoplasmata_A; RBG-16-68-12; RBG-16-68-12 | 88.8 | 2.4 | 2,235,584 | 75 | 2361 | MQ | 0 | 130.44 | 0.6 | 1,947,181 |
| East | 3300038455_25 | Chloroflexota | Bacteria; Chloroflexota; Ellin6529; CSP1-4; CSP1-4 | 55.06 | 1.94 | 1,946,312 | 306 | 2064 | MQ | 0 | 39.59 | 0.16 | 514,763 |
| East | 3300038455_27 | Proteobacteria | Bacteria; Proteobacteria; Alphaproteobacteria; Rhizobiales; Methyloligellaceae | 51.36 | 7.45 | 1,568,936 | 256 | 1776 | MQ | 0 | 13.23 | 0.04 | 138,647 |
| East | 3300038468_11 | Desulfobacterota | Bacteria; Desulfobacterota; Syntrophobacteria | 75.81 | 8.31 | 4,395,771 | 297 | 4376 | MQ | 1 | 19.70 | 0.22 | 578,771 |
| East | 3300038468_12 | UBP1 | Bacteria; UBP1; PRR-12; SSS58A | 85.51 | 5.49 | 3,537,679 | 406 | 3438 | MQ | 1 | 10.54 | 0.09 | 249,337 |
| East | 3300038468_13 | Nitrospirota | Bacteria; Nitrospirota; Nitrospiria; Nitrospirales; Nitrospiraceae; Nitrospira_C | 81.91 | 8.26 | 3,376,013 | 487 | 3936 | MQ | 1 | 22.48 | 0.19 | 507,254 |
| East | 3300038468_14 | Halobacterota | Archaea; Halobacterota; Methanosarcinia; Methanocellales; Methanocellaceae; Methanocella | 70.22 | 1.96 | 2,168,271 | 383 | 2624 | MQ | 0 | 23.94 | 0.13 | 347,320 |
| East | 3300038468_15 | Nitrospirota | Bacteria; Nitrospirota; Nitrospiria; 2-01-FULL-66-17 | 79.17 | 6.53 | 1,869,992 | 226 | 1964 | MQ | 0 | 12.70 | 0.06 | 158,786 |
| East | 3300038468_17 | Halobacterota | Archaea; Halobacterota; Methanosarcinia | 75.19 | 5.23 | 1,407,663 | 204 | 1638 | MQ | 0 | 8.83 | 0.03 | 83,074 |
| East | 3300038468_18 | Thermoplasmatota | Archaea; Thermoplasmatota; Thermoplasmata_A; RBG-16-68-12; RBG-16-68-12 | 66.53 | 1.26 | 1,250,379 | 198 | 1456 | MQ | 0 | 30.06 | 0.09 | 251,362 |
| East | 3300038468_21 | Nitrospirota | Bacteria; Nitrospirota; Thermodesulfovibrionia; Thermodesulfovibrionales; SM23-35 | 60.18 | 0.91 | 1,171,813 | 224 | 1341 | MQ | 0 | 9.55 | 0.03 | 74,794 |
| East | 3300038468_23 | Nitrospirota | Bacteria; Nitrospirota; Nitrospiria; 2-01-FULL-66-17 | 63.24 | 0.45 | 1,072,418 | 117 | 1184 | MQ | 0 | 9.01 | 0.02 | 64,573 |
| East | 3300038468_24 | Crenarchaeota | Archaea; Crenarchaeota; Bathyarchaeia; B26-1; BA1 | 56.03 | 6.21 | 947,501 | 211 | 1183 | MQ | 0 | 8.82 | 0.02 | 55,879 |
| North | 3300038409_13 | Chloroflexota | Bacteria; Chloroflexota; Anaerolineae; Anaerolineales | 56.82 | 5.91 | 3,434,748 | 549 | 3532 | MQ | 0 | 10.45 | 0.09 | 240,197 |
| North | 3300038409_18 | Chloroflexota | Bacteria; Chloroflexota; Ellin6529; CSP1-4; CSP1-4 | 55.13 | 0.93 | 2,049,605 | 292 | 2083 | MQ | 0 | 16.20 | 0.08 | 221,886 |
| North | 3300038409_21 | RBG-13-66-14 | Bacteria; RBG-13-66-14; RBG-13-66-14 | 57.87 | 1.23 | 1,400,574 | 267 | 1565 | MQ | 1 | 16.20 | 0.08 | 221,886 |
| North | 3300038409_9 | Acidobacteriota | Bacteria; Acidobacteriota; Acidobacteriae | 63.32 | 7 | 5,362,978 | 992 | 5462 | MQ | 2 | 9.95 | 0.13 | 356,820 |
| North | 3300038421_13 | Nitrospirota | Bacteria; Nitrospirota; Thermodesulfovibrionia; UBA6902 | 88.13 | 8.18 | 3,294,869 | 217 | 3588 | MQ | 1 | 18.21 | 0.14 | 401,138 |
| North | 3300038421_15 | Planctomycetota | Bacteria; Planctomycetota; Phycisphaerae; SG8-4; SG8-4; SG8-4 | 55.81 | 2.75 | 3,018,999 | 567 | 2956 | MQ | 1 | 10.56 | 0.07 | 213,166 |
| North | 3300038421_17 | Chloroflexota | Bacteria; Chloroflexota; Ellin6529; CSP1-4; CSP1-4 | 72.04 | 3.24 | 2,246,127 | 351 | 2311 | MQ | 0 | 17.46 | 0.09 | 262,178 |
| North | 3300038421_9 | Acidobacteriota | Bacteria; Acidobacteriota; Acidobacteriae | 78.5 | 4.7 | 6,949,535 | 1106 | 6950 | MQ | 2 | 10.34 | 0.16 | 480,397 |
| North | 3300038422_14 | Bacteroidota | Bacteria; Bacteroidota; Bacteroidia; Bacteroidales; SM23-62; SM23-62 | 68.13 | 3.73 | 3,637,539 | 629 | 3363 | MQ | 0 | 7.33 | 0.07 | 177,987 |
| North | 3300038422_19 | Desulfuromonadota | Bacteria; Desulfuromonadota; Desulfuromonadia; Geobacterales; Geobacteraceae | 53.28 | 2.58 | 1,787,540 | 320 | 1960 | MQ | 0 | 8.70 | 0.04 | 103,829 |
| North | 3300038422_22 | Proteobacteria | Bacteria; Proteobacteria; Gammaproteobacteria | 77.59 | 2.35 | 1,488,114 | 156 | 1634 | MQ | 1 | 14.02 | 0.06 | 139,225 |
| North | 3300038422_24 | Nitrospirota | Bacteria; Nitrospirota; Nitrospiria; 2-01-FULL-66-17 | 55.24 | 0.96 | 1,358,836 | 213 | 1548 | MQ | 0 | 8.93 | 0.03 | 81,055 |
| North | 3300038422_25 | Chloroflexota | Bacteria; Chloroflexota; Dehalococcoidia; Dehalococcoidales | 91.75 | 3.96 | 1,259,480 | 39 | 1363 | HQ | 1 | 18.36 | 0.06 | 155,778 |
| North | 3300038422_28 | Nitrospirota | Bacteria; Nitrospirota; Thermodesulfovibrionia; Thermodesulfovibrionales | 53.63 | 6.54 | 1,130,843 | 239 | 1346 | MQ | 0 | 11.07 | 0.03 | 83,577 |
| North | 3300038422_31 | Desulfobacterota | Bacteria; Desulfobacterota; BSN033; BSN033; RBG-16-54-18; RBG-13-52-11 | 52.17 | 1.18 | 958,529 | 194 | 1137 | MQ | 0 | 8.99 | 0.02 | 57,522 |
| North | 3300038550_24 | Thermoplasmatota | Archaea; Thermoplasmatota; E2; DHVEG-1; DHVEG-1; SM1-50 | 61.78 | 0 | 1,463,981 | 156 | 1619 | MQ | 1 | 14.30 | 0.05 | 139,817 |
| North | 3300038550_26 | Chloroflexota | Bacteria; Chloroflexota; Dehalococcoidia; Dehalococcoidales | 92.74 | 2.48 | 1,289,298 | 46 | 1399 | HQ | 2 | 23.20 | 0.07 | 199,914 |
| North | 3300038550_27 | Halobacterota | Archaea; Halobacterota; Methanosarcinia; Methanotrichales; Methanotrichaceae | 65.66 | 4.02 | 1,236,142 | 244 | 1441 | MQ | 0 | 7.99 | 0.02 | 65,972 |
| North | 3300038550_28 | MBNT15 | Bacteria; MBNT15; MBNT15; MBNT15; MBNT15; RBG-16-64-85 | 52.4 | 1.68 | 1,199,162 | 164 | 1330 | MQ | 0 | 10.96 | 0.03 | 87,837 |
| North | 3300038550_30 | Chloroflexota | Bacteria; Chloroflexota; Dehalococcoidia; Dehalococcoidales; RBG-16-60-22; RBG-16-56-11 | 56.2 | 1.98 | 1,146,258 | 226 | 1349 | MQ | 0 | 16.72 | 0.04 | 128,114 |
| North | 3300038550_31 | Proteobacteria | Bacteria; Proteobacteria; Gammaproteobacteria | 52.69 | 0.35 | 1,102,817 | 156 | 1226 | MQ | 0 | 21.03 | 0.05 | 154,883 |
| North | 3300038550_32 | Acidobacteriota | Bacteria; Acidobacteriota; Aminicenantia; Aminicenantales; Aminicenantaceae | 54.3 | 0.85 | 1,092,575 | 234 | 1192 | MQ | 0 | 8.59 | 0.02 | 62,707 |
| North | 3300038558_17 | Acidobacteriota | Bacteria; Acidobacteriota; Acidobacteriae | 59.69 | 0.85 | 4,257,154 | 827 | 4309 | MQ | 0 | 11.48 | 0.09 | 326,427 |
| North | 3300038558_20 | Desulfuromonadota | Bacteria; Desulfuromonadota; Desulfuromonadia; Geobacterales; Geobacteraceae | 64.04 | 1.75 | 3,182,487 | 398 | 3451 | MQ | 0 | 15.64 | 0.09 | 332,346 |
| North | 3300038558_21 | Bacteroidota | Bacteria; Bacteroidota; Ignavibacteria; Ignavibacteriales; Ignavibacteriaceae; RBG-16-36-9 | 68.9 | 2.44 | 2,655,382 | 437 | 2717 | MQ | 0 | 9.52 | 0.05 | 168,663 |
| North | 3300038558_23 | Zixibacteria | Bacteria; Zixibacteria; MSB-5A5; UBA10806; UBA10806 | 70.5 | 1.1 | 2,256,919 | 405 | 2330 | MQ | 0 | 9.11 | 0.04 | 137,333 |
| North | 3300038558_25 | Halobacterota | Archaea; Halobacterota; Methanosarcinia; Methanotrichales; Methanotrichaceae | 92.16 | 0.98 | 2,017,734 | 223 | 2248 | MQ | 1 | 11.69 | 0.04 | 157,416 |
| North | 3300038558_27 | Thermoplasmatota | Archaea; Thermoplasmatota; E2; DHVEG-1; DHVEG-1; SM1-50 | 67.81 | 0.8 | 1,626,184 | 254 | 1770 | MQ | 1 | 26.42 | 0.08 | 286,924 |
| North | 3300038558_29 | Desulfobacterota | Bacteria; Desulfobacterota; Desulfobaccia; Desulfobaccales; 0-14-0-80-60-11; 0-14-0-80-60-11 | 51.82 | 0.65 | 1,699,637 | 341 | 1843 | MQ | 0 | 8.96 | 0.03 | 101,827 |
| North | 3300038558_31 | Chloroflexota | Bacteria; Chloroflexota; Dehalococcoidia; Dehalococcoidales | 90.76 | 0.99 | 1,244,817 | 35 | 1345 | HQ | 1 | 24.01 | 0.06 | 199,469 |
| North | 3300038558_32 | Proteobacteria | Bacteria; Proteobacteria; Gammaproteobacteria | 61.03 | 0.79 | 1,235,436 | 206 | 1440 | MQ | 1 | 8.85 | 0.02 | 72,981 |
| North | 3300038558_36 | Chloroflexota | Bacteria; Chloroflexota; Dehalococcoidia; Dehalococcoidales; RBG-16-60-22; RBG-16-56-11 | 67.43 | 0.99 | 1,125,109 | 211 | 1314 | MQ | 0 | 16.68 | 0.04 | 125,259 |
| North | 3300038558_39 | Crenarchaeota | Archaea; Crenarchaeota; Methanomethylicia; Methanomethyliales; Methanomethyliaceae | 79.44 | 0.93 | 928,103 | 100 | 1146 | MQ | 1 | 9.71 | 0.02 | 60,216 |
| North | 3300038558_40 | Chloroflexota | Bacteria; Chloroflexota; Dehalococcoidia; Dehalococcoidales; UBA2162 | 58.75 | 3.96 | 955,367 | 176 | 1084 | MQ | 1 | 14.08 | 0.03 | 89,808 |
| North | 3300038558_42 | Halobacterota | Archaea; Halobacterota; Methanomicrobia; Methanomicrobiales; Methanofollaceae | 51.96 | 2.29 | 859,578 | 171 | 1031 | MQ | 0 | 8.11 | 0.01 | 46,518 |

**Supplementary Table S1, continued.**

| Site | Bin ID | Phylum | Taxonomy | Completeness | Contamination | Genome Size (bp) | Scaffold Count | Gene Count | MIMAG classification | 16S | Mean Coverage | Mapped Reads (%) | Number of Mapped Reads |
|---|---|---|---|---|---|---|---|---|---|---|---|---|---|
| South | 3300038401_2 | Proteobacteria | Bacteria; Proteobacteria; Gammaproteobacteria; Beggiatoales; Beggiatoaceae | 74.17 | 0.88 | 3,135,568 | 498 | 2861 | MQ | 1 | 8.87 | 0.08 | 182,470 |
| South | 3300038403_10 | Thermoplasmatota | Archaea; Thermoplasmatota; Thermoplasmata_A; RBG-16-68-12; RBG-16-68-12 | 53.78 | 1.6 | 825,276 | 103 | 957 | MQ | 1 | 13.59 | 0.03 | 74,927 |
| South | 3300038549_10 | Thermoplasmatota | Archaea; Thermoplasmatota; Thermoplasmata_A; UBA10834; UBA10834; RBG-16-62-10 | 79.01 | 3.64 | 1,491,661 | 224 | 1672 | MQ | 0 | 14.70 | 0.04 | 146,508 |
| South | 3300038549_11 | Proteobacteria | Bacteria; Proteobacteria; Gammaproteobacteria; Betaproteobacteriales; Hydrogenophilaceae; UBA3361 | 50.41 | 2.53 | 1,741,428 | 373 | 2037 | MQ | 2 | 14.72 | 0.05 | 171,365 |
| South | 3300038549_15 | Thermoplasmatota | Archaea; Thermoplasmatota; Thermoplasmata_A; RBG-16-68-12; RBG-16-68-12; COMBO-69-17 | 52.93 | 1.6 | 821,559 | 114 | 938 | MQ | 0 | 18.50 | 0.03 | 101,557 |
| South | 3300038549_3 | Acidobacteriota | Bacteria; Acidobacteriota; Blastocatellia; Pyrinomonadales; Pyrinomonadaceae | 97.44 | 3.42 | 7,339,912 | 152 | 6252 | HQ | 1 | 16.94 | 0.25 | 830,431 |
| South | 3300038663_10 | Asgardarchaeota | Archaea; Asgardarchaeota; Lokiarchaeia | 88.79 | 1.87 | 2,082,377 | 278 | 2308 | MQ | 1 | 9.87 | 0.05 | 137,294 |
| South | 3300038663_17 | Nitrospirota | Bacteria; Nitrospirota; Thermodesulfovibrionia; Thermodesulfovibrionales; SM23-35 | 56.44 | 2.4 | 1,413,628 | 250 | 1615 | MQ | 2 | 11.22 | 0.04 | 106,065 |
| West | 3300038408_13 | Acidobacteriota | Bacteria; Acidobacteriota; Blastocatellia; Pyrinomonadales | 78.56 | 4.27 | 4,528,148 | 714 | 4047 | MQ | 1 | 8.82 | 0.08 | 266,929 |
| West | 3300038408_17 | Chloroflexota | Bacteria; Chloroflexota; Anaerolineae; 4572-78 | 53.54 | 6.09 | 3,109,501 | 632 | 3218 | MQ | 0 | 9.68 | 0.06 | 201,127 |
| West | 3300038408_18 | Zixibacteria | Bacteria; Zixibacteria; MSB-5A5; UBA10806; UBA10806; UBA10806 | 71.82 | 1.1 | 2,827,498 | 332 | 2694 | MQ | 0 | 10.21 | 0.06 | 192,894 |
| West | 3300038408_20 | Proteobacteria | Bacteria; Proteobacteria; Gammaproteobacteria | 67.25 | 1.56 | 2,823,466 | 520 | 3189 | MQ | 18 | 15.34 | 0.09 | 289,648 |
| West | 3300038408_29 | Desulfobacterota | Bacteria; Desulfobacterota; BSN033; BSN033; RBG-16-54-18; RBG-13-52-11 | 82.9 | 3.35 | 2,018,850 | 145 | 2168 | MQ | 1 | 16.08 | 0.06 | 216,937 |
| West | 3300038408_30 | Chloroflexota | Bacteria; Chloroflexota; Dehalococcoidia; GIF9; AB-539-J10; RBG-13-51-36 | 71.48 | 9.16 | 1,939,560 | 269 | 2097 | MQ | 1 | 13.69 | 0.05 | 177,418 |
| West | 3300038408_32 | Bacteroidota | Bacteria; Bacteroidota; Ignavibacteria; SJA-28; OLB5 | 51.34 | 0.56 | 1,845,501 | 329 | 1868 | MQ | 0 | 9.91 | 0.04 | 122,330 |
| West | 3300038408_35 | Nitrospirota | Bacteria; Nitrospirota; Thermodesulfovibrionia; Thermodesulfovibrionales; JdFR-86 | 71.67 | 0.97 | 1,627,111 | 269 | 1914 | MQ | 1 | 11.66 | 0.04 | 126,833 |
| West | 3300038408_36 | Thermoplasmatota | Archaea; Thermoplasmatota; Thermoplasmata_A; RBG-16-68-12; RBG-16-68-12 | 64.46 | 0.8 | 1,268,108 | 131 | 1453 | MQ | 1 | 14.76 | 0.04 | 125,084 |
| West | 3300038408_37 | Nitrospirota | Bacteria; Nitrospirota; Thermodesulfovibrionia; Thermodesulfovibrionales; SM23-35 | 51.75 | 1.75 | 1,233,434 | 197 | 1403 | MQ | 0 | 11.54 | 0.03 | 95,143 |
| West | 3300038408_40 | Halobacterota | Archaea; Halobacterota; Methanomicrobia; Methanomicrobiales | 72.61 | 1.31 | 1,115,752 | 201 | 1364 | MQ | 0 | 7.67 | 0.02 | 57,218 |
| West | 3300038408_46 | Crenarchaeota | Archaea; Crenarchaeota; Nitrososphaeria; Nitrososphaerales; Nitrosopumilaceae; CSP1-1 | 55.35 | 0.49 | 565,187 | 115 | 789 | MQ | 0 | 17.65 | 0.02 | 66,690 |
| West | 3300038469_28 | Firmicutes | Bacteria; Firmicutes; Bacilli; Bacillales; Bacillaceae_G; Bacillus_A; Bacillus_A cereus_E | 73.62 | 0.2 | 3,564,137 | 113 | 3851 | MQ | 1 | 11.63 | 0.06 | 277,631 |
| West | 3300038469_29 | Calditrichota | Bacteria; Calditrichota; Calditrichia; Calditrichales | 75.8 | 4.4 | 2,741,665 | 485 | 2699 | MQ | 1 | 8.67 | 0.04 | 158,966 |
| West | 3300038469_32 | Nitrospirota | Bacteria; Nitrospirota; Thermodesulfovibrionia; Thermodesulfovibrionales | 72.3 | 6.57 | 2,411,196 | 285 | 2671 | MQ | 0 | 15.05 | 0.05 | 242,515 |
| West | 3300038469_33 | Zixibacteria | Bacteria; Zixibacteria; MSB-5A5; UBA10806; UBA10806; UBA10806 | 66.3 | 3.4 | 2,319,569 | 395 | 2306 | MQ | 1 | 8.14 | 0.03 | 126,453 |
| West | 3300038469_34 | Halobacterota | Archaea; Halobacterota; Methanomicrobia; Methanomicrobiales | 85.13 | 4.58 | 1,926,313 | 222 | 2334 | MQ | 2 | 18.53 | 0.05 | 238,549 |
| West | 3300038469_36 | Desulfobacterota | Bacteria; Desulfobacterota; BSN033; BSN033; RBG-16-54-18; RBG-13-52-11 | 82.29 | 4.19 | 1,588,412 | 124 | 1748 | MQ | 1 | 20.58 | 0.05 | 218,630 |
| West | 3300038469_37 | Nitrospirota | Bacteria; Nitrospirota; Thermodesulfovibrionia; Thermodesulfovibrionales; SM23-35 | 56.57 | 5.45 | 1,490,733 | 278 | 1704 | MQ | 1 | 10.69 | 0.02 | 106,469 |
| West | 3300038469_39 | Halobacterota | Archaea; Halobacterota; Methanomicrobia; Methanomicrobiales; Methanoregulaceae; Methanoregula | 57.69 | 6.86 | 1,270,306 | 248 | 1593 | MQ | 1 | 11.54 | 0.02 | 97,956 |
| West | 3300038469_40 | Chloroflexota | Bacteria; Chloroflexota; Dehalococcoidia; Dehalococcoidales; RBG-16-60-22; RBG-13-51-18 | 57.62 | 5.94 | 1,200,482 | 258 | 1429 | MQ | 1 | 9.53 | 0.02 | 76,459 |
| West | 3300038469_41 | Crenarchaeota | Archaea; Crenarchaeota; Bathyarchaeia; 40CM-2-53-6; RBG-16-50-20 | 62.4 | 4.69 | 1,046,328 | 134 | 1260 | MQ | 0 | 18.16 | 0.03 | 126,948 |
| West | 3300038470_17 | Firmicutes | Bacteria; Firmicutes; Bacilli; Bacillales; Bacillaceae_G; Bacillus_A; Bacillus_A cereus_E | 85.25 | 2.52 | 4,260,441 | 442 | 4815 | MQ | 2 | 8.67 | 0.07 | 247,038 |
| West | 3300038470_18 | Bacteroidota | Bacteria; Bacteroidota; Ignavibacteria; Ignavibacteriales; Ignavibacteriaceae | 78.4 | 2.12 | 4,085,017 | 634 | 4107 | MQ | 1 | 9.29 | 0.07 | 253,393 |
| West | 3300038470_19 | Chloroflexota | Bacteria; Chloroflexota; Anaerolineae; 4572-78 | 61.21 | 8.15 | 4,106,905 | 808 | 4282 | MQ | 0 | 10.55 | 0.08 | 289,528 |
| West | 3300038470_29 | Chloroflexota | Bacteria; Chloroflexota; Dehalococcoidia; GIF9; AB-539-J10; RBG-13-51-36 | 70.9 | 3.96 | 1,615,195 | 232 | 1735 | MQ | 3 | 20.00 | 0.06 | 215,798 |
| West | 3300038470_30 | Halobacterota | Archaea; Halobacterota; Methanomicrobia; Methanomicrobiales | 83.13 | 9.44 | 1,572,354 | 246 | 1984 | MQ | 1 | 8.78 | 0.03 | 92,193 |
| West | 3300038470_34 | Desulfobacterota | Bacteria; Desulfobacterota; BSN033; BSN033; RBG-16-54-18; RBG-13-52-11 | 59.39 | 0 | 1,044,287 | 180 | 1228 | MQ | 0 | 8.88 | 0.02 | 61,974 |
| West | 3300038551_24 | Chloroflexota | Bacteria; Chloroflexota; Anaerolineae; 4572-78 | 64.29 | 7.27 | 4,618,128 | 847 | 4709 | MQ | 1 | 11.34 | 0.08 | 350,230 |
| West | 3300038551_29 | Proteobacteria | Bacteria; Proteobacteria; Gammaproteobacteria | 74.84 | 1.76 | 3,543,491 | 507 | 3952 | MQ | 30 | 17.73 | 0.1 | 420,230 |
| West | 3300038551_36 | Zixibacteria | Bacteria; Zixibacteria; MSB-5A5; UBA10806; UBA10806; UBA10806 | 72.53 | 1.1 | 2,294,965 | 387 | 2275 | MQ | 1 | 8.22 | 0.03 | 126,229 |
| West | 3300038551_39 | Calditrichota | Bacteria; Calditrichota; Calditrichia; Calditrichales | 54.76 | 0 | 1,958,813 | 394 | 1951 | MQ | 1 | 7.47 | 0.02 | 97,721 |
| West | 3300038551_40 | Halobacterota | Archaea; Halobacterota; Methanomicrobia; Methanomicrobiales | 91.45 | 6.8 | 1,867,625 | 175 | 2253 | MQ | 1 | 20.25 | 0.06 | 246,471 |
| West | 3300038551_41 | Desulfobacterota | Bacteria; Desulfobacterota; BSN033; BSN033; RBG-16-54-18; RBG-13-52-11 | 81.68 | 2.9 | 1,812,936 | 167 | 2014 | MQ | 0 | 17.09 | 0.05 | 207,198 |
| West | 3300038551_43 | Eisenbacteria | Bacteria; Eisenbacteria; RBG-16-71-46 | 50.55 | 1.1 | 1,147,036 | 233 | 1168 | MQ | 0 | 13.41 | 0.04 | 176,495 |
| West | 3300038551_49 | Crenarchaeota | Archaea; Crenarchaeota; Bathyarchaeia; B26-1; UBA233; AD8-1 | 57.87 | 3.74 | 924,895 | 190 | 1157 | MQ | 0 | 10.96 | 0.02 | 67,779 |
| West | 3300038551_50 | Methylomirabilota | Bacteria; Methylomirabilota; Methylomirabilia | 51.15 | 1.72 | 798,177 | 142 | 874 | MQ | 0 | 10.70 | 0.01 | 57,047 |
| West | 3300038552_17 | Desulfobacterota | Bacteria; Desulfobacterota; Desulfobacteria; Desulfobacterales; Desulfosarcinaceae | 98.71 | 5.44 | 6,817,238 | 189 | 6595 | MQ | 1 | 15.88 | 0.15 | 722,715 |
| West | 3300038552_26 | Proteobacteria | Bacteria; Proteobacteria; Gammaproteobacteria | 76.98 | 2.97 | 3,364,301 | 540 | 3724 | MQ | 3 | 11.48 | 0.05 | 258,244 |
| West | 3300038552_28 | Actinobacteriota | Bacteria; Actinobacteriota; Actinobacteria; Actinomycetales; Kineosporiaceae | 63.83 | 9.61 | 3,286,775 | 570 | 3032 | MQ | 0 | 9.42 | 0.04 | 206,934 |
| West | 3300038552_30 | Zixibacteria | Bacteria; Zixibacteria; MSB-5A5; UBA10806; UBA10806; UBA10806 | 74.88 | 1.78 | 2,712,718 | 409 | 2673 | MQ | 0 | 9.29 | 0.04 | 168,412 |
| West | 3300038552_31 | Calditrichota | Bacteria; Calditrichota; Calditrichia; Calditrichales | 72.54 | 1.1 | 2,323,398 | 432 | 2289 | MQ | 0 | 7.65 | 0.02 | 118,621 |
| West | 3300038552_35 | Nitrospirota | Bacteria; Nitrospirota; Thermodesulfovibrionia; Thermodesulfovibrionales; UBA1546 | 72.48 | 4.33 | 1,970,022 | 233 | 2215 | MQ | 1 | 14.99 | 0.04 | 197,220 |
| West | 3300038552_36 | Desulfobacterota | Bacteria; Desulfobacterota; BSN033; BSN033; RBG-16-54-18; RBG-13-52-11 | 83.94 | 3.42 | 1,820,730 | 175 | 2023 | MQ | 1 | 17.47 | 0.04 | 212,472 |
| West | 3300038552_37 | Nitrospirota | Bacteria; Nitrospirota; Thermodesulfovibrionia; Thermodesulfovibrionales; SM23-35 | 69.79 | 4.55 | 1,804,619 | 250 | 2016 | MQ | 0 | 17.79 | 0.04 | 214,385 |
| West | 3300038552_38 | Halobacterota | Archaea; Halobacterota; Methanomicrobia; Methanomicrobiales | 85.62 | 2.88 | 1,820,695 | 193 | 2216 | MQ | 1 | 17.61 | 0.04 | 212,140 |
| West | 3300038552_48 | Crenarchaeota | Archaea; Crenarchaeota; Bathyarchaeia; B26-1; UBA233; 20-14-0-80-47-9 | 54.67 | 1.89 | 879,257 | 140 | 1043 | MQ | 0 | 19.74 | 0.02 | 115,921 |
| West | 3300038552_51 | Crenarchaeota | Archaea; Crenarchaeota; Bathyarchaeia; 40CM-2-53-6; RBG-16-50-20 | 56.63 | 0 | 750,560 | 120 | 918 | MQ | 0 | 10.51 | 0.01 | 52,681 |

**Supplementary Table S2:** Pearson's correlation of water (aq) and sediment analytes versus estimated gene frequencies, as determined by IMG pipeline.

| Analyte | | Gene: *nifH* | *narG* | *phoD* | *phnX* | *kdpB* | *kup* | *dsrA* | *cysC* | *cbbL* | *mcrA* |
|---|---|---|---|---|---|---|---|---|---|---|---|
| pH (aq) | p | 0.4900 | 0.2110 | 0.0030 | 0.0000 | 0.9700 | 0.0020 | 0.0030 | 0.1000 | 0.0140 | 0.1840 |
| | *R* | 0.1640 | 0.2930 | 0.6330 | -0.7660 | 0.0090 | 0.6590 | 0.6220 | -0.3780 | 0.5400 | 0.3090 |
| TN (aq) | p | 0.3190 | 0.4040 | 0.3440 | 0.0170 | 0.0140 | 0.5690 | 0.0070 | 0.4950 | 0.1840 | 0.5450 |
| | *R* | -0.2350 | -0.1980 | 0.2230 | -0.5260 | -0.5390 | 0.1360 | 0.5840 | 0.1620 | 0.3100 | -0.1440 |
| TOC (aq) | p | 0.0320 | 0.0980 | 0.9040 | 0.0180 | 0.0010 | 0.6880 | 0.0460 | 0.2390 | 0.8170 | 0.0850 |
| | *R* | -0.4810 | -0.3800 | 0.0290 | -0.5230 | -0.6780 | -0.0960 | 0.4510 | 0.2760 | 0.0550 | -0.3950 |
| K (aq) | p | 0.0000 | 0.0040 | 0.1900 | 0.1420 | 0.0000 | 0.0380 | 0.5160 | 0.0470 | 0.1520 | 0.0010 |
| | *R* | -0.7490 | -0.6090 | -0.3050 | -0.3400 | -0.7820 | -0.4670 | 0.1540 | 0.4500 | -0.3320 | -0.7010 |
| Mg (aq) | p | 0.0000 | 0.0050 | 0.2210 | 0.1250 | 0.0000 | 0.0490 | 0.4620 | 0.0510 | 0.1840 | 0.0010 |
| | *R* | -0.7370 | -0.5980 | -0.2860 | -0.3540 | -0.7810 | -0.4460 | 0.1740 | 0.4420 | -0.3100 | -0.6860 |
| Ammonia (aq) | p | 0.0000 | 0.0010 | 0.0010 | 0.6000 | 0.0090 | 0.0000 | 0.0990 | 0.0220 | 0.0000 | 0.0000 |
| | *R* | 0.7950 | 0.6720 | 0.6780 | -0.1250 | 0.5690 | 0.8220 | 0.3790 | -0.5100 | 0.7650 | 0.8340 |
| Nitrate (aq) | p | 0.6960 | 0.4110 | 0.6720 | 0.1010 | 0.1340 | 0.5130 | 0.3580 | 0.0520 | 0.2180 | 0.9340 |
| | *R* | 0.0930 | -0.1940 | -0.1010 | 0.3780 | -0.3470 | -0.1550 | 0.2170 | 0.4410 | 0.2880 | 0.0200 |
| EC (aq) | p | 0.0030 | 0.0020 | 0.0520 | 0.9130 | 0.0000 | 0.0050 | 0.8230 | 0.0040 | 0.1840 | 0.0020 |
| | *R* | -0.6330 | -0.6440 | -0.4400 | 0.0260 | -0.7720 | -0.6000 | 0.0530 | 0.6150 | -0.3090 | -0.6530 |
| Cl (aq) | p | 0.5390 | 0.4380 | 0.2510 | 0.0010 | 0.4140 | 0.1950 | 0.8340 | 0.0390 | 0.6090 | 0.9350 |
| | *R* | 0.1460 | -0.1840 | -0.2690 | 0.6720 | -0.1930 | -0.3020 | -0.0500 | 0.4640 | 0.1220 | 0.0190 |
| S (aq) | p | 0.0020 | 0.0050 | 0.1870 | 0.4360 | 0.0000 | 0.0380 | 0.4230 | 0.0170 | 0.3510 | 0.0040 |
| | *R* | -0.6380 | -0.5980 | -0.3070 | -0.1840 | -0.7910 | -0.4660 | 0.1900 | 0.5280 | -0.2200 | -0.6190 |
| Ortho-P (aq) | p | 0.4550 | 0.0730 | 0.0090 | 0.0000 | 0.1930 | 0.0020 | 0.1600 | 0.0070 | 0.2070 | 0.1700 |
| | *R* | 0.1770 | 0.4090 | 0.5710 | -0.7300 | 0.3040 | 0.6430 | 0.3260 | -0.5850 | 0.2950 | 0.3190 |
| Ca (aq) | p | 0.0000 | 0.0010 | 0.0040 | 0.5130 | 0.0000 | 0.0000 | 0.4670 | 0.0030 | 0.0140 | 0.0000 |
| | *R* | -0.7200 | -0.7010 | -0.6100 | 0.1550 | -0.7170 | -0.7680 | -0.1730 | 0.6370 | -0.5400 | -0.7650 |
| pH (sediment) | p | 0.9270 | 0.5110 | 0.0420 | 0.0160 | 0.1500 | 0.0650 | 0.0200 | 0.6470 | 0.0070 | 0.5970 |
| | *R* | -0.0230 | 0.1660 | 0.4840 | -0.5580 | -0.3540 | 0.4450 | 0.5430 | 0.1160 | 0.6120 | 0.1340 |
| %TN (sediment) | p | 0.2641 | 0.0062 | 0.1135 | 0.1949 | 0.0087 | 0.0450 | 0.4118 | 0.0019 | 0.8908 | 0.1840 |
| | *R* | -0.2622 | -0.5894 | -0.3651 | 0.3025 | -0.5698 | -0.4528 | 0.1943 | 0.6504 | 0.0328 | -0.3096 |
| %TOC (sediment) | p | 0.0889 | 0.0068 | 0.0755 | 0.3787 | 0.0024 | 0.0207 | 0.4879 | 0.0011 | 0.7564 | 0.0641 |
| | *R* | -0.3903 | -0.5845 | -0.4063 | 0.2081 | -0.6394 | -0.5130 | 0.1646 | 0.6761 | -0.0740 | -0.4217 |

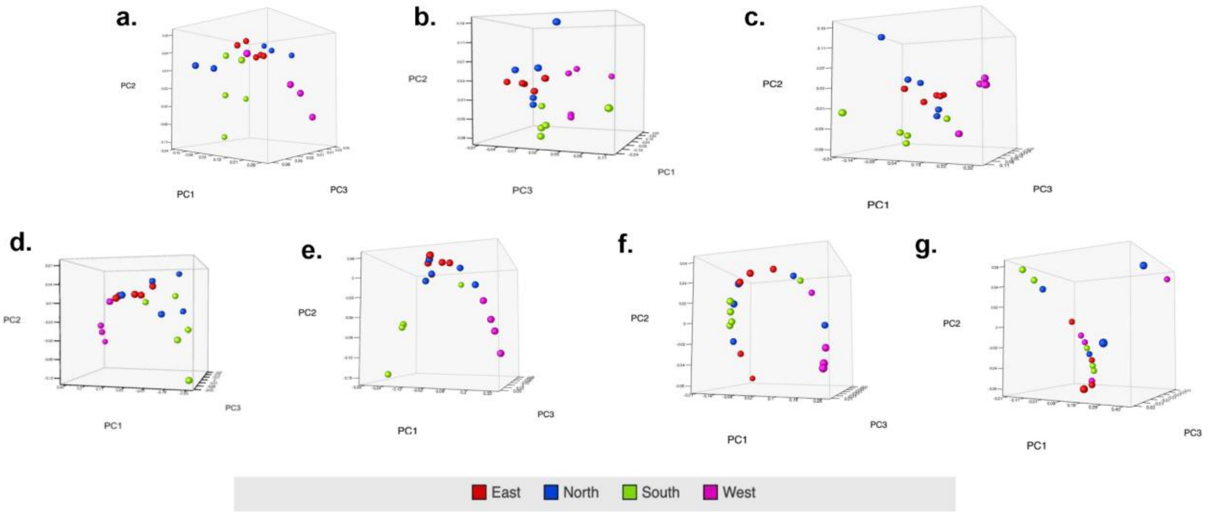

**Supplementary Figure S3:** PCoA analysis of each functional gene group. a: Overall (all 10 KOs utilized in this study); b: *nifH/narG*; c: *phoD/phnX*; d: *kdpB/kup*; e: *dsrA/cysC*; f: *cbbL*; g: *mcrA.*

**Supplementary Table S3:** Summary of high quality bins mapped against sequenced read from each metagenomic sample within their respective sites.

| Bin ID | Metagenomic Sample | Mapped Against | Mapped Reads (%) | Number of Mapped Reads |
|---|---|---|---|---|
| 3300038454_18 | East 1 | East 1 | 0.06 | 169,697 |
| 3300038454_18 | East 1 | East 2 | 0.07 | 219,039 |
| 3300038454_18 | East 1 | East 3 | 0.02 | 71,336 |
| 3300038454_18 | East 1 | East 4 | 0.07 | 165,523 |
| 3300038454_18 | East 1 | East 5 | 0.05 | 130,060 |
| 3300038454_19 | East 1 | East 1 | 0.36 | 1,085,636 |
| 3300038454_19 | East 1 | East 2 | 0.53 | 1,720,931 |
| 3300038454_19 | East 1 | East 3 | 0.25 | 743,074 |
| 3300038454_19 | East 1 | East 4 | 0.33 | 819,097 |
| 3300038454_19 | East 1 | East 5 | 0.15 | 404,842 |
| 3300038558_31 | North 3 | North 1 | 0.01 | 20,516 |
| 3300038558_31 | North 3 | North 2 | 0 | 12,909 |
| 3300038558_31 | North 3 | North 3 | 0.06 | 199,469 |
| 3300038558_31 | North 3 | North 4 | 0.07 | 197,072 |
| 3300038558_31 | North 3 | North 5 | 0.06 | 154,166 |
| 3300038550_26 | North 4 | North 1 | 0.01 | 19,986 |
| 3300038550_26 | North 4 | North 2 | 0 | 12,223 |
| 3300038550_26 | North 4 | North 3 | 0.06 | 203,176 |
| 3300038550_26 | North 4 | North 4 | 0.07 | 199,914 |
| 3300038550_26 | North 4 | North 5 | 0.07 | 156,405 |
| 3300038422_25 | North 5 | North 1 | 0.01 | 20,651 |
| 3300038422_25 | North 5 | North 2 | 0 | 12,968 |
| 3300038422_25 | North 5 | North 3 | 0.06 | 201,372 |
| 3300038422_25 | North 5 | North 4 | 0.07 | 198,920 |
| 3300038422_25 | North 5 | North 5 | 0.06 | 155,778 |
| 3300038549_3 | South 2 | South 1 | 0 | 8,456 |
| 3300038549_3 | South 2 | South 2 | 0.25 | 830,431 |
| 3300038549_3 | South 2 | South 3 | 0.01 | 14,694 |
| 3300038549_3 | South 2 | South 4 | 0.01 | 35,896 |
| 3300038549_3 | South 2 | South 5 | 0.02 | 54,469 |

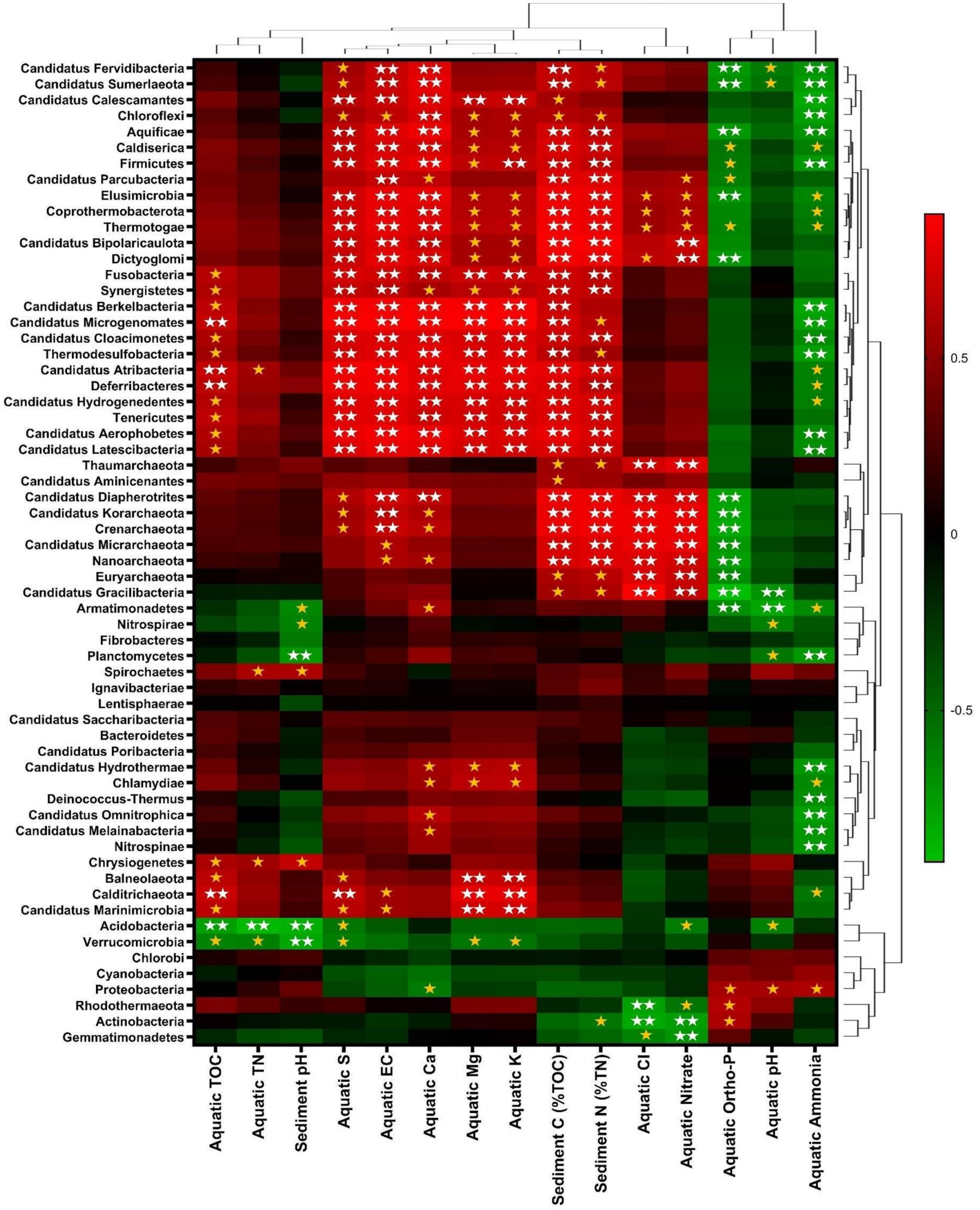

**Supplementary Figure S4:** Heatmap of Pearson's correlation of analytes versus estimated gene frequencies of phyla. Heatmap colors indicate *R* value, single gold star = *p* <.01, double-white star = *p* <.001.

**Supplementary Table S4:** Genes and metabolic pathways analyzed in this research.

| Gene | KO | Protein Products | Metabolic Pathway/KO |
|---|---|---|---|
| ***nifH*** | **K02588** | Nitrogenase iron protein | Nitrogen metabolism (KO00910) |
| ***narG*** | **K00370** | Nitrate reductase / nitrite oxidoreductase, alpha subunit | Nitrogen metabolism (KO00910) |
| ***cbbL*** | **K01601** | Ribulose-bisphosphate carboxylase large chain | Carbohydrate metabolism (KO09101) |
| ***phoD*** | **K01113** | Alkaline Phosphatase | Hydrolyzing organic phosphates (KO01100) |
| ***phnX*** | **K05306** | Phosphonoacetaldehyde hydrolase | Hydrolysis of organic phosphates (KO01100) |
| ***kdpB*** | **K01547** | Potassium-transporting ATPase ATP-binding subunit | Potassium cycling/transport (KO01000) |
| ***kup*** | **K03549** | KUP system potassium uptake protein | Electrochemical potential-driven transporters (KO02000) |
| ***dsrA*** | **K11180** | Dissimilatory Sulfate Reductase alpha subunit | Sulfur metabolism (KO00920) |
| ***cysC*** | **K00860** | Adenylyl-sulfate kinase | Sulfur metabolism (KO00920) |
| ***mcrA*** | **K00399** | Methyl-coenzyme M reductase alpha subunit | Methane metabolism (KO00680) |

**Supplementary Table S5:** Additional frequencies of archaea found within metagenomic samples which are not included in Figure 2d.

| Site | Unclassified Archaeoglobales | Halobacteriaceae | Haloferacaceae | Halorubraceae | Unclassified Haloferacales | Methano-bacteriaceae | Methanocellaceae | Methanomass-iliicoccaceae | Unclassified Methano-massiliicoccales | Methanocorp-usculaceae | Methanomicro-biaceae | Methano-spirillaceae | Unclassified Methano-microbiales | Candidatus Methano-peredenaceae | Methano-sarcinaceae | Methano-trichaceae | Unclassified Methanosarcinales | Natrialbaceae |
|---|---|---|---|---|---|---|---|---|---|---|---|---|---|---|---|---|---|---|
| North 1 | 7.36E-02 | 5.65E-02 | 7.25E-02 | 6.56E-02 | 7.36E-02 | 6.12E-02 | 5.91E-02 | 4.74E-02 | 7.36E-02 | 5.24E-02 | 2.70E-02 | 5.71E-02 | 7.36E-02 | 5.09E-02 | 7.56E-02 | 4.33E-02 | 7.36E-02 | 7.63E-02 |
| North 2 | 7.84E-02 | 5.94E-02 | 7.85E-02 | 7.45E-02 | 7.84E-02 | 1.01E-01 | 6.32E-02 | 5.17E-02 | 7.84E-02 | 5.63E-02 | 3.01E-02 | 6.23E-02 | 7.84E-02 | 5.16E-02 | 8.18E-02 | 5.00E-02 | 7.84E-02 | 7.61E-02 |
| North 3 | 1.63E-01 | 9.56E-02 | 1.06E-01 | 1.16E-01 | 1.63E-01 | 1.86E-01 | 1.15E-01 | 1.52E-01 | 1.63E-01 | 1.60E-01 | 1.63E-01 | 1.48E-01 | 1.63E-01 | 8.01E-02 | 1.49E-01 | 1.39E-01 | 1.63E-01 | 1.04E-01 |
| North 4 | 1.45E-01 | 7.93E-02 | 9.64E-02 | 1.02E-01 | 1.45E-01 | 1.57E-01 | 9.63E-02 | 1.44E-01 | 1.45E-01 | 1.54E-01 | 1.47E-01 | 1.29E-01 | 1.45E-01 | 6.88E-02 | 1.26E-01 | 1.11E-01 | 1.45E-01 | 9.03E-02 |
| North 5 | 1.03E-01 | 5.67E-02 | 7.13E-02 | 7.13E-02 | 1.03E-01 | 9.45E-02 | 7.16E-02 | 1.27E-01 | 1.03E-01 | 9.94E-02 | 1.19E-01 | 9.12E-02 | 1.03E-01 | 4.55E-02 | 8.45E-02 | 9.20E-02 | 1.03E-01 | 6.59E-02 |
| South 1 | 7.27E-03 | 6.34E-03 | 1.00E-02 | 9.89E-03 | 7.27E-03 | 1.09E-02 | 4.08E-03 | 7.85E-03 | 7.27E-03 | 2.28E-02 | 1.99E-02 | 2.02E-02 | 7.27E-03 | 3.15E-03 | 9.45E-03 | 2.53E-02 | 7.27E-03 | 1.03E-02 |
| South 2 | 2.46E-02 | 2.53E-02 | 3.71E-02 | 3.77E-02 | 2.46E-02 | 2.96E-02 | 1.52E-02 | 3.90E-02 | 2.46E-02 | 5.65E-02 | 6.03E-02 | 5.48E-02 | 2.46E-02 | 1.33E-02 | 2.86E-02 | 1.03E-01 | 2.46E-02 | 4.03E-02 |
| South 3 | 1.02E-02 | 8.45E-03 | 1.48E-02 | 1.61E-02 | 1.02E-02 | 4.17E-02 | 8.18E-03 | 9.74E-03 | 1.02E-02 | 4.20E-02 | 5.58E-02 | 4.58E-02 | 1.02E-02 | 6.50E-03 | 1.92E-02 | 8.61E-02 | 1.02E-02 | 1.51E-02 |
| South 4 | 1.66E-02 | 1.46E-02 | 2.40E-02 | 2.35E-02 | 1.66E-02 | 1.56E-02 | 1.26E-02 | 1.84E-02 | 1.66E-02 | 3.31E-02 | 4.54E-02 | 3.08E-02 | 1.66E-02 | 1.00E-02 | 1.94E-02 | 2.53E-02 | 1.66E-02 | 2.39E-02 |
| South 5 | 3.47E-02 | 2.81E-02 | 3.91E-02 | 4.37E-02 | 3.47E-02 | 3.28E-02 | 2.74E-02 | 4.17E-02 | 3.47E-02 | 3.87E-02 | 6.71E-02 | 4.70E-02 | 3.47E-02 | 2.27E-02 | 3.90E-02 | 3.82E-02 | 3.47E-02 | 4.04E-02 |
| West 1 | 8.32E-03 | 1.01E-02 | 1.43E-02 | 1.56E-02 | 8.32E-03 | 9.36E-03 | 5.70E-03 | 4.00E-03 | 8.32E-03 | 9.69E-03 | 6.48E-03 | 1.10E-02 | 8.32E-03 | 5.71E-03 | 1.07E-02 | 9.92E-03 | 8.32E-03 | 1.46E-02 |
| West 2 | 6.21E-03 | 7.46E-03 | 1.16E-02 | 1.16E-02 | 6.21E-03 | 6.78E-03 | 4.31E-03 | 2.71E-03 | 6.21E-03 | 5.64E-03 | 4.26E-03 | 7.40E-03 | 6.21E-03 | 4.47E-03 | 7.94E-03 | 5.05E-03 | 6.21E-03 | 1.25E-02 |
| West 3 | 1.04E-02 | 1.27E-02 | 1.80E-02 | 1.90E-02 | 1.04E-02 | 1.20E-02 | 7.25E-03 | 4.82E-03 | 1.04E-02 | 1.16E-02 | 8.00E-03 | 1.42E-02 | 1.04E-02 | 7.23E-03 | 1.38E-02 | 1.97E-02 | 1.04E-02 | 1.88E-02 |
| West 4 | 1.09E-02 | 1.18E-02 | 1.60E-02 | 1.68E-02 | 1.09E-02 | 1.23E-02 | 7.32E-03 | 5.01E-03 | 1.09E-02 | 1.38E-02 | 8.60E-03 | 1.32E-02 | 1.09E-02 | 7.14E-03 | 1.35E-02 | 1.12E-02 | 1.09E-02 | 1.59E-02 |
| West 5 | 6.93E-03 | 7.57E-03 | 1.01E-02 | 1.12E-02 | 6.93E-03 | 7.93E-03 | 4.64E-03 | 3.12E-03 | 6.93E-03 | 7.48E-03 | 4.86E-03 | 7.83E-03 | 6.93E-03 | 4.67E-03 | 8.57E-03 | 5.58E-03 | 6.93E-03 | 1.11E-02 |
| East 1 | 7.02E-02 | 5.94E-02 | 8.79E-02 | 8.13E-02 | 7.02E-02 | 5.21E-02 | 1.07E-01 | 7.96E-02 | 7.02E-02 | 6.06E-02 | 5.69E-02 | 6.21E-02 | 7.02E-02 | 1.53E-01 | 7.36E-02 | 5.79E-02 | 7.02E-02 | 9.11E-02 |
| East 2 | 8.76E-02 | 6.22E-02 | 9.67E-02 | 9.71E-02 | 8.76E-02 | 6.20E-02 | 9.20E-02 | 1.03E-01 | 8.76E-02 | 6.11E-02 | 6.46E-02 | 7.18E-02 | 8.76E-02 | 2.04E-01 | 8.74E-02 | 6.24E-02 | 8.76E-02 | 9.77E-02 |
| East 3 | 4.85E-02 | 4.10E-02 | 6.73E-02 | 6.40E-02 | 4.85E-02 | 3.30E-02 | 4.47E-02 | 4.83E-02 | 4.85E-02 | 4.72E-02 | 5.15E-02 | 4.94E-02 | 4.85E-02 | 9.52E-02 | 5.08E-02 | 4.42E-02 | 4.85E-02 | 6.59E-02 |
| East 4 | 5.42E-02 | 4.24E-02 | 6.93E-02 | 6.75E-02 | 5.42E-02 | 3.94E-02 | 8.63E-02 | 6.04E-02 | 5.42E-02 | 3.66E-02 | 3.15E-02 | 3.92E-02 | 5.42E-02 | 9.89E-02 | 5.31E-02 | 3.17E-02 | 5.42E-02 | 7.28E-02 |
| East 5 | 4.03E-02 | 3.74E-02 | 5.87E-02 | 5.56E-02 | 4.03E-02 | 3.41E-02 | 1.68E-01 | 4.99E-02 | 4.03E-02 | 3.09E-02 | 2.87E-02 | 3.82E-02 | 4.03E-02 | 6.71E-02 | 4.76E-02 | 3.84E-02 | 4.03E-02 | 5.74E-02 |

**Supplementary Table S6a:** Actual and mean values of water analytes. BDL= below detectable limits (<0.005 mg/L).

| Water Analysis | Analyte | | | | | | | | | | | | | | | |
|---|---|---|---|---|---|---|---|---|---|---|---|---|---|---|---|---|
| Sample | pH | TN (mg/L) | TOC (mg/L) | P (mg/L) | K (mg/L) | Mg (mg/L) | Fe (mg/L) | Mn (mg/L) | Ammonia_N (mg/L) | Nitrate-N (mg/L) | Pb (mg/L0 | EC (dS/m) | Cl (mg/L) | S (mg/L) | Ortho-P (ug/L) | Ca (mg/L) |
| SOUTH 1 | 8.01 | 0.74 | 53.56 | 0.040 | 2.95 | 5.39 | BDL | BDL | 0.16 | 0.03 | BDL | 0.34 | 46.85 | 3.99 | 4.27 | 25.91 |
| SOUTH 2 | 8.2 | 0.79 | 61.93 | 0.010 | 2.11 | 3.83 | BDL | BDL | 0.06 | 0.02 | BDL | 0.24 | 30.77 | 2.82 | 3.97 | 18.32 |
| SOUTH 3 | 8.35 | 0.71 | 51.55 | 0.010 | 2.94 | 4.98 | BDL | BDL | 0.05 | 0.01 | BDL | 0.32 | 46.33 | 3.65 | 3.54 | 23.97 |
| NORTH 1 | 7.55 | 0.99 | 71.75 | 0.010 | 3.28 | 6.07 | BDL | BDL | 0.04 | 0.03 | BDL | 0.41 | 56.42 | 5.82 | 3.53 | 32.23 |
| NORTH 2 | 7.98 | 0.67 | 51.11 | 0.010 | 5.19 | 9.34 | BDL | BDL | 0.05 | 0.05 | BDL | 0.57 | 87.10 | 8.74 | 3.69 | 30.72 |
| NORTH 3 | 7.55 | 0.70 | 62.46 | BDL | 3.56 | 6.15 | BDL | BDL | 0.05 | 0.03 | BDL | 0.42 | 56.98 | 5.81 | 4.02 | 31.57 |
| EAST 1 | 7.67 | 0.54 | 52.17 | 0.010 | 2.53 | 4.48 | BDL | BDL | 0.04 | 0.01 | BDL | 0.30 | 39.89 | 3.26 | 3.95 | 24.99 |
| EAST 2 | 7.51 | 0.46 | 45.22 | 0.010 | 3.00 | 5.54 | BDL | BDL | 0.05 | 0.01 | BDL | 0.35 | 30.68 | 3.90 | 3.86 | 28.42 |
| EAST 3 | 7.55 | 0.49 | 45.73 | BDL | 3.40 | 5.53 | BDL | BDL | 0.05 | 0.01 | BDL | 0.37 | 51.93 | 3.81 | 3.72 | 29.66 |
| WEST 1 | 8.09 | 0.48 | 47.15 | BDL | 5.14 | 9.07 | BDL | BDL | 0.04 | 0.01 | BDL | 0.48 | 21.47 | 7.49 | 3.73 | 33.87 |
| WEST 2 | 7.97 | 0.81 | 66.16 | BDL | 4.61 | 8.87 | BDL | BDL | 0.04 | 0.03 | BDL | 0.44 | 61.72 | 7.32 | 3.88 | 28.07 |
| WEST 3 | 8.27 | 1.38 | 96.14 | BDL | 4.75 | 8.32 | BDL | BDL | 0.05 | 0.01 | BDL | 0.44 | 20.75 | 6.81 | 4.09 | 29.95 |
| **Mean Values** | | | | | | | | | | | | | | | | |
| South | 8.19 | 0.75 | 55.68 | 0.02 | 2.67 | 4.73 | BDL | BDL | 0.09 | 0.02 | BDL | 0.30 | 41.31 | 3.49 | 3.93 | 22.73 |
| North | 7.69 | 0.79 | 61.77 | 0.01 | 4.01 | 7.19 | BDL | BDL | 0.05 | 0.04 | BDL | 0.47 | 66.83 | 6.79 | 3.75 | 31.51 |
| East | 7.58 | 0.49 | 47.71 | 0.01 | 2.97 | 5.18 | BDL | BDL | 0.05 | 0.01 | BDL | 0.34 | 40.84 | 3.66 | 3.84 | 27.69 |
| West | 8.11 | 0.89 | 69.82 | BDL | 4.83 | 8.75 | BDL | BDL | 0.04 | 0.02 | BDL | 0.46 | 34.64 | 7.21 | 3.90 | 30.63 |

**Supplementary Table S6b:** Actual and mean values of sediment analytes. * = Low biomass in the sample could have impacted the accuracy of these values. ** = Not enough biomass in the samples for any analysis.

| Sediment Analysis | | Analyte | | |
|---|---|---|---|---|
| | Sample | pH | Carbon (%TOC) | Nitrogen (%TN) |
| | SOUTH 1 | ** | 7.69 | 0.94 |
| | SOUTH 2 | ** | 3.37 | 0.09 |
| | SOUTH 3 | 7.85 | 2.18 | 0.04 |
| | SOUTH 4 | 7.86 | 2.40 | 0.02 |
| | SOUTH 5 | 7.86 | 2.97 | 0.02 |
| | NORTH 1 | 7.54* | 40.64 | 2.66 |
| | NORTH 2 | 7.61 | 31.63 | 1.62 |
| | NORTH 3 | 7.50* | 43.85 | 3.03 |
| | NORTH 4 | 7.46* | 45.24 | 2.93 |
| | NORTH 5 | 7.33* | 46.02 | 3.25 |
| | EAST 1 | 6.53 | 5.75 | 0.27 |
| | EAST 2 | 6.59 | 4.15 | 0.15 |
| | EAST 3 | 6.81 | 11.51 | 0.54 |
| | EAST 4 | 6.59 | 2.69 | 0.05 |
| | EAST 5 | 5.61 | 6.28 | 0.37 |
| | WEST 1 | 7.70 | 26.25 | 1.47 |
| | WEST 2 | 7.60 | 27.36 | 1.58 |
| | WEST 3 | 7.68 | 26.90 | 1.53 |
| | WEST 4 | 7.66 | 25.31 | 1.31 |
| | WEST 5 | 7.69 | 23.89 | 1.25 |
| | **Mean Values** | | | |
| | South | 7.86 | 3.72 | 0.22 |
| | North | 7.49 | 41.48 | 2.70 |
| | East | 6.43 | 6.08 | 0.28 |
| | West | 7.67 | 25.94 | 1.43 |

**Supplementary Table S7:** Metadata from each cardinal boundary the day of collection.

| Sample Site | Water Temp (ºC) | Water Depth (m) | Air Temp (ºC) | Wind Speed (m/s) |
|---|---|---|---|---|
| North | n/a* | 0.457 | 16.2 | 1.1 |
| South | 22.5 | 0.432 | 24.7 | 1.3 |
| East | n/a* | 0.457 | 20.4 | 1.2 |
| West | 21.0 | 0.838 | 19.7 | 1.3 |

*Aquatic thermometer broken in the field.

**Supplementary Table S8:** Table of the estimated raw gene counts of each gene per site versus the normalized gene frequencies per site, as determined by IMG pipeline.

| | Sample | Measurement | Gene | | | | | | | | | |
|---|---|---|---|---|---|---|---|---|---|---|---|---|
| | | | *nifH* | *narG* | *phoD* | *phnX* | *kdpB* | *kup* | *dsrA* | *cysC* | *cbbL* | *mcrA* |
| Genome | Lox_East_1 | Estimated Raw Gene Count | 70 | 486 | 248 | 63 | 1,086 | 386 | 139 | 170 | 225 | 28 |
| | | Normalized Gene Frequency | 6.39E-05 | 4.44E-04 | 2.26E-04 | 5.75E-05 | 9.92E-04 | 3.52E-04 | 1.27E-04 | 1.55E-04 | 2.05E-04 | 2.56E-05 |
| | Lox_East_2 | Estimated Raw Gene Count | 77 | 504 | 257 | 53 | 1,158 | 356 | 161 | 172 | 253 | 31 |
| | | Normalized Gene Frequency | 6.83E-05 | 4.47E-04 | 2.28E-04 | 4.70E-05 | 1.03E-03 | 3.16E-04 | 1.43E-04 | 1.52E-04 | 2.24E-04 | 2.75E-05 |
| | Lox_East_3 | Estimated Raw Gene Count | 58 | 446 | 227 | 47 | 953 | 369 | 152 | 155 | 212 | 25 |
| | | Normalized Gene Frequency | 5.98E-05 | 4.60E-04 | 2.34E-04 | 4.85E-05 | 9.83E-04 | 3.80E-04 | 1.57E-04 | 1.60E-04 | 2.19E-04 | 2.58E-05 |
| | Lox_East_4 | Estimated Raw Gene Count | 52 | 448 | 245 | 54 | 945 | 312 | 132 | 141 | 166 | 16 |
| | | Normalized Gene Frequency | 5.59E-05 | 4.81E-04 | 2.63E-04 | 5.80E-05 | 1.02E-03 | 3.35E-04 | 1.42E-04 | 1.52E-04 | 1.78E-04 | 1.72E-05 |
| | Lox_East_5 | Estimated Raw Gene Count | 53 | 367 | 192 | 47 | 916 | 284 | 134 | 114 | 153 | 20 |
| | | Normalized Gene Frequency | 6.15E-05 | 4.26E-04 | 2.23E-04 | 5.46E-05 | 1.06E-03 | 3.30E-04 | 1.56E-04 | 1.32E-04 | 1.78E-04 | 2.32E-05 |
| | Lox_North_1 | Estimated Raw Gene Count | 39 | 362 | 260 | 42 | 644 | 275 | 110 | 120 | 172 | 12 |
| | | Normalized Gene Frequency | 4.96E-05 | 4.60E-04 | 3.30E-04 | 5.34E-05 | 8.18E-04 | 3.49E-04 | 1.40E-04 | 1.52E-04 | 2.19E-04 | 1.52E-05 |
| | Lox_North_2 | Estimated Raw Gene Count | 47 | 366 | 263 | 45 | 719 | 346 | 130 | 122 | 194 | 9 |
| | | Normalized Gene Frequency | 5.43E-05 | 4.23E-04 | 3.04E-04 | 5.20E-05 | 8.31E-04 | 4.00E-04 | 1.50E-04 | 1.41E-04 | 2.24E-04 | 1.04E-05 |
| | Lox_North_3 | Estimated Raw Gene Count | 90 | 498 | 200 | 67 | 1,063 | 374 | 224 | 215 | 377 | 35 |
| | | Normalized Gene Frequency | 7.81E-05 | 4.32E-04 | 1.74E-04 | 5.82E-05 | 9.23E-04 | 3.25E-04 | 1.94E-04 | 1.87E-04 | 3.27E-04 | 3.04E-05 |
| | Lox_North_4 | Estimated Raw Gene Count | 67 | 362 | 191 | 52 | 929 | 276 | 178 | 180 | 282 | 26 |
| | | Normalized Gene Frequency | 7.11E-05 | 3.84E-04 | 2.03E-04 | 5.52E-05 | 9.86E-04 | 2.93E-04 | 1.89E-04 | 1.91E-04 | 2.99E-04 | 2.76E-05 |
| | Lox_North_5 | Estimated Raw Gene Count | 53 | 294 | 144 | 51 | 759 | 221 | 136 | 150 | 210 | 28 |
| | | Normalized Gene Frequency | 7.03E-05 | 3.90E-04 | 1.91E-04 | 6.77E-05 | 1.01E-03 | 2.93E-04 | 1.80E-04 | 1.99E-04 | 2.79E-04 | 3.72E-05 |
| | Lox_South_1 | Estimated Raw Gene Count | 62 | 175 | 212 | 23 | 427 | 302 | 84 | 47 | 195 | 24 |
| | | Normalized Gene Frequency | 1.38E-04 | 3.89E-04 | 4.71E-04 | 5.11E-05 | 9.49E-04 | 6.71E-04 | 1.87E-04 | 1.04E-04 | 4.33E-04 | 5.34E-05 |
| | Lox_South_2 | Estimated Raw Gene Count | 91 | 462 | 296 | 42 | 910 | 470 | 171 | 120 | 295 | 36 |
| | | Normalized Gene Frequency | 1.07E-04 | 5.44E-04 | 3.49E-04 | 4.95E-05 | 1.07E-03 | 5.54E-04 | 2.01E-04 | 1.41E-04 | 3.48E-04 | 4.24E-05 |
| | Lox_South_3 | Estimated Raw Gene Count | 66 | 261 | 157 | 17 | 570 | 308 | 106 | 62 | 188 | 37 |
| | | Normalized Gene Frequency | 1.28E-04 | 5.06E-04 | 3.05E-04 | 3.30E-05 | 1.11E-03 | 5.97E-04 | 2.06E-04 | 1.20E-04 | 3.65E-04 | 7.18E-05 |
| | Lox_South_4 | Estimated Raw Gene Count | 54 | 311 | 213 | 17 | 564 | 283 | 93 | 72 | 180 | 27 |
| | | Normalized Gene Frequency | 9.74E-05 | 5.61E-04 | 3.84E-04 | 3.07E-05 | 1.02E-03 | 5.10E-04 | 1.68E-04 | 1.30E-04 | 3.25E-04 | 4.87E-05 |
| | Lox_South_5 | Estimated Raw Gene Count | 38 | 418 | 203 | 21 | 732 | 270 | 118 | 112 | 183 | 23 |
| | | Normalized Gene Frequency | 5.63E-05 | 6.19E-04 | 3.01E-04 | 3.11E-05 | 1.08E-03 | 4.00E-04 | 1.75E-04 | 1.66E-04 | 2.71E-04 | 3.41E-05 |
| | Lox_West_1 | Estimated Raw Gene Count | 51 | 660 | 490 | 54 | 1,533 | 644 | 284 | 270 | 406 | 21 |
| | | Normalized Gene Frequency | 2.95E-05 | 3.81E-04 | 2.83E-04 | 3.12E-05 | 8.86E-04 | 3.72E-04 | 1.64E-04 | 1.56E-04 | 2.35E-04 | 1.21E-05 |
| | Lox_West_2 | Estimated Raw Gene Count | 38 | 623 | 367 | 37 | 1,201 | 475 | 285 | 209 | 322 | 9 |
| | | Normalized Gene Frequency | 2.81E-05 | 4.61E-04 | 2.71E-04 | 2.74E-05 | 8.88E-04 | 3.51E-04 | 2.11E-04 | 1.55E-04 | 2.38E-04 | 6.66E-06 |
| | Lox_West_3 | Estimated Raw Gene Count | 60 | 846 | 566 | 47 | 1,798 | 742 | 342 | 316 | 427 | 29 |
| | | Normalized Gene Frequency | 2.96E-05 | 4.17E-04 | 2.79E-04 | 2.32E-05 | 8.86E-04 | 3.66E-04 | 1.69E-04 | 1.56E-04 | 2.10E-04 | 1.43E-05 |
| | Lox_West_4 | Estimated Raw Gene Count | 57 | 674 | 452 | 53 | 1,631 | 683 | 299 | 291 | 422 | 28 |
| | | Normalized Gene Frequency | 3.19E-05 | 3.77E-04 | 2.53E-04 | 2.97E-05 | 9.13E-04 | 3.82E-04 | 1.67E-04 | 1.63E-04 | 2.36E-04 | 1.57E-05 |
| | Lox_West_5 | Estimated Raw Gene Count | 37 | 543 | 372 | 38 | 1,111 | 518 | 249 | 199 | 324 | 17 |
| | | Normalized Gene Frequency | 2.84E-05 | 4.16E-04 | 2.85E-04 | 2.91E-05 | 8.52E-04 | 3.97E-04 | 1.91E-04 | 1.53E-04 | 2.48E-04 | 1.30E-05 |

**Supplementary Table S9:** Raw Spearman's correlation of water and sediment analytes versus alpha diversity results. **(a)** Bacterial analysis. **(b)** Archaeal analysis.

| a. | | | Bacterial Diversity Analysis | | | |
|---|---|---|---|---|---|---|
| | | | Margalef index | Simpson index D | Shannon index | Pielou index |
| Analyte | pH (aq) | *p* value | 0.183 | 0.002 | 0.006 | 0.006 |
| | | Rho (ρ) | 0.310 | -0.651 | -0.589 | -0.589 |
| | TN (aq) | *p* value | 0.023 | 0.745 | 0.948 | 0.948 |
| | | Rho (ρ) | -0.504 | -0.078 | 0.016 | 0.016 |
| | TOC (aq) | *p* value | 0.023 | 0.745 | 0.948 | 0.948 |
| | | Rho (ρ) | -0.504 | -0.078 | 0.016 | 0.016 |
| | K (aq) | *p* value | 0.000 | 0.161 | 0.091 | 0.091 |
| | | Rho (ρ) | -0.822 | 0.326 | 0.388 | 0.388 |
| | Mg (aq) | *p* value | 0.000 | 0.161 | 0.091 | 0.091 |
| | | Rho (ρ) | -0.822 | 0.326 | 0.388 | 0.388 |
| | Ammonia (aq) | *p* value | 0.000 | 0.067 | 0.041 | 0.041 |
| | | Rho (ρ) | 0.897 | -0.417 | -0.461 | -0.461 |
| | Nitrate (aq) | *p* value | 0.078 | 0.340 | 0.413 | 0.413 |
| | | Rho (ρ) | 0.403 | -0.225 | -0.194 | -0.194 |
| | EC (aq) | *p* value | 0.029 | 0.066 | 0.047 | 0.047 |
| | | Rho (ρ) | -0.489 | 0.419 | 0.450 | 0.450 |
| | Cl (aq) | *p* value | 0.002 | 0.795 | 0.696 | 0.696 |
| | | Rho (ρ) | 0.659 | -0.062 | -0.093 | -0.093 |
| | S (aq) | *p* value | 0.000 | 0.161 | 0.091 | 0.091 |
| | | Rho (ρ) | -0.822 | 0.326 | 0.388 | 0.388 |
| | Ortho-P (aq) | *p* value | 0.324 | 0.007 | 0.012 | 0.012 |
| | | Rho (ρ) | 0.233 | -0.582 | -0.551 | -0.551 |
| | Ca (aq) | *p* value | 0.029 | 0.066 | 0.047 | 0.047 |
| | | Rho (ρ) | -0.489 | 0.419 | 0.450 | 0.450 |
| | pH (sediment) | *p* value | 0.871 | 0.002 | 0.008 | 0.008 |
| | | Rho (ρ) | 0.041 | -0.675 | -0.606 | -0.606 |
| | %TN (sediment) | *p* value | 0.112 | 0.146 | 0.099 | 0.099 |
| | | Rho (ρ) | -0.371 | 0.337 | 0.379 | 0.379 |
| | %TOC (sediment) | *p* value | 0.107 | 0.121 | 0.081 | 0.081 |
| | | Rho (ρ) | -0.371 | 0.358 | 0.400 | 0.400 |

| b. | | | Archaeal Diversity Analysis | | | |
|---|---|---|---|---|---|---|
| | | | Margalef index | Simpson index D | Shannon index | Pielou index |
| Analyte | pH (aq) | *p* value | 0.672 | 0.721 | 0.922 | 0.922 |
| | | Rho (ρ) | 0.101 | 0.085 | -0.023 | -0.023 |
| | TN (aq) | *p* value | 0.023 | 0.013 | 0.002 | 0.002 |
| | | Rho (ρ) | -0.504 | 0.543 | 0.651 | 0.651 |
| | TOC (aq) | *p* value | 0.023 | 0.013 | 0.002 | 0.002 |
| | | Rho (ρ) | -0.504 | 0.543 | 0.651 | 0.651 |
| | K (aq) | *p* value | 0.015 | 0.072 | 0.004 | 0.004 |
| | | Rho (ρ) | -0.535 | 0.411 | 0.613 | 0.613 |
| | Mg (aq) | *p* value | 0.015 | 0.072 | 0.004 | 0.004 |
| | | Rho (ρ) | -0.535 | 0.411 | 0.613 | 0.613 |
| | Ammonia (aq) | *p* value | 0.269 | 0.997 | 0.351 | 0.351 |
| | | Rho (ρ) | 0.260 | -0.001 | -0.220 | -0.220 |
| | Nitrate (aq) | *p* value | 0.114 | 0.002 | 0.010 | 0.010 |
| | | Rho (ρ) | -0.365 | 0.651 | 0.558 | 0.558 |
| | EC (aq) | *p* value | 0.002 | 0.003 | 0.000 | 0.000 |
| | | Rho (ρ) | -0.644 | 0.636 | 0.776 | 0.776 |
| | Cl (aq) | *p* value | 0.535 | 0.078 | 0.261 | 0.261 |
| | | Rho (ρ) | -0.147 | 0.403 | 0.264 | 0.264 |
| | S (aq) | *p* value | 0.015 | 0.072 | 0.004 | 0.004 |
| | | Rho (ρ) | -0.535 | 0.411 | 0.613 | 0.613 |
| | Ortho-P (aq) | *p* value | 0.061 | 0.091 | 0.032 | 0.032 |
| | | Rho (ρ) | 0.427 | -0.388 | -0.481 | -0.481 |
| | Ca (aq) | *p* value | 0.002 | 0.003 | 0.000 | 0.000 |
| | | Rho (ρ) | -0.644 | 0.636 | 0.776 | 0.776 |
| | pH (sediment) | *p* value | 0.523 | 0.191 | 0.319 | 0.319 |
| | | Rho (ρ) | -0.161 | 0.323 | 0.249 | 0.249 |
| | %TN (sediment) | *p* value | 0.012 | 0.026 | 0.003 | 0.003 |
| | | Rho (ρ) | -0.551 | 0.496 | 0.627 | 0.627 |
| | %TOC (sediment) | *p* value | 0.022 | 0.012 | 0.001 | 0.001 |
| | | Rho (ρ) | -0.508 | 0.552 | 0.669 | 0.669 |

**Supplementary Table S10:** Calculated alpha diversity raw values.

| | Bacterial Phyla | | | | | | | | Archaeal Phyla | | | | | | | |
|---|---|---|---|---|---|---|---|---|---|---|---|---|---|---|---|---|
| | Shannon's Index | | Simpson's Index | | Margalef's Richness | | Pielou's Evenness | | Shannon's Index | | Simpson's Index | | Margalef's Richness | | Pielou's Evenness | |
| Sample | IMG | KAIJU | IMG | KAIJU | IMG | KAIJU | IMG | KAIJU | IMG | KAIJU | IMG | KAIJU | IMG | KAIJU | IMG | KAIJU |
| Lox_East_1 | 1.848993 | 1.818244 | 0.669048 | 0.667483 | 3.203931 | 2.374005 | 0.461402 | 0.499849 | 0.652663 | 0.613529 | 0.325413 | 0.303507 | 0.436245 | 0.319371 | 0.335403 | 0.381207 |
| Lox_East_2 | 1.877078 | 1.856462 | 0.682951 | 0.681753 | 3.200735 | 2.363815 | 0.468411 | 0.510356 | 0.683954 | 0.624259 | 0.347309 | 0.309786 | 0.430737 | 0.317272 | 0.351483 | 0.387874 |
| Lox_East_3 | 1.770672 | 1.814122 | 0.639906 | 0.664237 | 3.236908 | 2.375224 | 0.441858 | 0.498716 | 0.613968 | 0.564846 | 0.301849 | 0.273383 | 0.44874 | 0.320685 | 0.315517 | 0.350959 |
| Lox_East_4 | 1.807902 | 1.804015 | 0.661492 | 0.66488 | 3.246563 | 2.402531 | 0.451148 | 0.495937 | 0.703527 | 0.6528 | 0.356898 | 0.327396 | 0.446841 | 0.325942 | 0.361541 | 0.405608 |
| Lox_East_5 | 1.848908 | 1.842622 | 0.666806 | 0.676823 | 3.267949 | 2.398744 | 0.461381 | 0.506551 | 0.645089 | 0.596655 | 0.320785 | 0.292713 | 0.449034 | 0.324391 | 0.33151 | 0.370723 |
| Lox_North_1 | 1.56914 | 1.639288 | 0.556745 | 0.59744 | 3.244971 | 2.389607 | 0.391567 | 0.450653 | 1.007134 | 0.81682 | 0.554879 | 0.432225 | 0.429397 | 0.322966 | 0.517564 | 0.507519 |
| Lox_North_2 | 1.585439 | 1.653961 | 0.561563 | 0.603042 | 3.224158 | 2.373726 | 0.395635 | 0.454686 | 0.997162 | 0.800549 | 0.548663 | 0.422285 | 0.427869 | 0.32106 | 0.51244 | 0.497409 |
| Lox_North_3 | 1.935968 | 1.881515 | 0.685779 | 0.685246 | 3.233762 | 2.358414 | 0.483106 | 0.517243 | 0.934571 | 0.756515 | 0.505471 | 0.393003 | 0.407682 | 0.306603 | 0.480274 | 0.470049 |
| Lox_North_4 | 1.988058 | 1.905873 | 0.704255 | 0.693809 | 3.275831 | 2.389327 | 0.496105 | 0.523939 | 0.930576 | 0.761446 | 0.503504 | 0.395421 | 0.411777 | 0.30928 | 0.478222 | 0.473113 |
| Lox_North_5 | 1.95023 | 1.900998 | 0.696392 | 0.693883 | 3.320215 | 2.419416 | 0.486665 | 0.522599 | 0.865152 | 0.715324 | 0.462401 | 0.365927 | 0.42373 | 0.3162 | 0.4446 | 0.444456 |
| Lox_South_1 | 1.433284 | 1.847375 | 0.511128 | 0.681948 | 3.443966 | 2.408905 | 0.357665 | 0.507857 | 0.355432 | 0.449041 | 0.157889 | 0.204596 | 0.498071 | 0.335292 | 0.182656 | 0.279005 |
| Lox_South_2 | 1.606656 | 1.831265 | 0.568658 | 0.672076 | 3.302813 | 2.358403 | 0.400929 | 0.503429 | 0.675314 | 0.618339 | 0.389095 | 0.313854 | 0.449156 | 0.318438 | 0.347043 | 0.384195 |
| Lox_South_3 | 1.466441 | 1.824681 | 0.51341 | 0.669495 | 3.418883 | 2.411918 | 0.365939 | 0.501619 | 0.54435 | 0.540144 | 0.301361 | 0.261082 | 0.388388 | 0.323389 | 0.303807 | 0.33561 |
| Lox_South_4 | 1.589605 | 1.85151 | 0.56495 | 0.681228 | 3.394673 | 2.407575 | 0.396674 | 0.508994 | 0.818705 | 0.684671 | 0.487198 | 0.357311 | 0.465001 | 0.326525 | 0.420731 | 0.42541 |
| Lox_South_5 | 1.719469 | 1.835196 | 0.610722 | 0.672561 | 3.344581 | 2.403839 | 0.429081 | 0.504509 | 0.868417 | 0.760237 | 0.491732 | 0.402726 | 0.447888 | 0.321584 | 0.446278 | 0.472362 |
| Lox_West_1 | 1.79639 | 1.755178 | 0.636358 | 0.643428 | 3.109028 | 2.319169 | 0.448276 | 0.482512 | 0.832566 | 0.706054 | 0.441659 | 0.360339 | 0.42798 | 0.313431 | 0.427854 | 0.438696 |
| Lox_West_2 | 1.7899 | 1.769242 | 0.632744 | 0.645772 | 3.163346 | 2.356149 | 0.446656 | 0.486378 | 0.882263 | 0.764513 | 0.477443 | 0.398447 | 0.43888 | 0.320728 | 0.453393 | 0.475018 |
| Lox_West_3 | 1.842397 | 1.777425 | 0.653287 | 0.651887 | 3.081558 | 2.306282 | 0.459756 | 0.488627 | 0.844291 | 0.735693 | 0.451558 | 0.379402 | 0.419485 | 0.309715 | 0.43388 | 0.457112 |
| Lox_West_4 | 1.82419 | 1.761231 | 0.646835 | 0.645611 | 3.102093 | 2.315361 | 0.455213 | 0.484176 | 0.865492 | 0.714993 | 0.460985 | 0.366297 | 0.41747 | 0.309616 | 0.444775 | 0.44425 |
| Lox_West_5 | 1.759007 | 1.730406 | 0.621789 | 0.634142 | 3.174131 | 2.349064 | 0.438947 | 0.475702 | 0.882988 | 0.711934 | 0.470232 | 0.364152 | 0.433788 | 0.318235 | 0.453766 | 0.44235 |

| | Prokaryotic Genera | | | |
|---|---|---|---|---|
| Sample | Shannon's Index | Simpson's Index | Margalef's Richness | Pielou's Evenness |
| Lox_East_1 | 5.343451 | 0.992727 | 19.97552 | 0.933585 |
| Lox_East_2 | 5.347726 | 0.992806 | 19.89371 | 0.934332 |
| Lox_East_3 | 5.356173 | 0.992967 | 19.92981 | 0.936343 |
| Lox_East_4 | 5.314707 | 0.992375 | 20.18993 | 0.928563 |
| Lox_East_5 | 5.340951 | 0.992708 | 20.18932 | 0.933148 |
| Lox_North_1 | 5.341855 | 0.992463 | 20.12371 | 0.933306 |
| Lox_North_2 | 5.346095 | 0.992503 | 19.99009 | 0.934047 |
| Lox_North_3 | 5.411914 | 0.993665 | 19.85625 | 0.945546 |
| Lox_North_4 | 5.40485 | 0.993618 | 20.11332 | 0.944312 |
| Lox_North_5 | 5.397737 | 0.993521 | 20.3706 | 0.943069 |
| Lox_South_1 | 5.44794 | 0.994391 | 20.33325 | 0.951841 |
| Lox_South_2 | 5.435846 | 0.994125 | 19.86472 | 0.949727 |
| Lox_South_3 | 5.423344 | 0.99396 | 20.31378 | 0.947543 |
| Lox_South_4 | 5.436347 | 0.994126 | 20.29165 | 0.949815 |
| Lox_South_5 | 5.399069 | 0.993628 | 20.17177 | 0.943842 |
| Lox_West_1 | 5.401423 | 0.993776 | 19.54287 | 0.943713 |
| Lox_West_2 | 5.388942 | 0.993541 | 19.86145 | 0.941533 |
| Lox_West_3 | 5.39011 | 0.99361 | 19.425 | 0.941737 |
| Lox_West_4 | 5.402922 | 0.99379 | 19.50451 | 0.943975 |
| Lox_West_5 | 5.414973 | 0.993911 | 19.80397 | 0.946081 |
